# CD8 memory precursor cells generation is a continuous process

**DOI:** 10.1101/2022.02.09.479673

**Authors:** Helena Todorov, Margaux Prieux, Daphne Laubreton, Matteo Bouvier, Shaoying Wang, Simon de Bernard, Christophe Arpin, Robrecht Cannoodt, Wouter Saelens, Arnaud Bonnaffoux, Olivier Gandrillon, Fabien Crauste, Yvan Saeys, Jacqueline Marvel

## Abstract

In this work, we studied the generation of memory precursor cells following an acute infection by analyzing single-cell RNA-seq data that contained CD8 T cells collected during the postinfection expansion phase. We used different tools to reconstruct the developmental trajectory that CD8 T cells followed after activation. Cells that exhibited a memory precursor signature were identified and positioned on this trajectory. We found that memory precursors are generated continuously with increasing numbers being generated over time. Similarly, expression of genes associated with effector functions was also found to be raised in memory precursors at later time points. The ability of cells to enter quiescence to differentiate into memory cells was confirmed by BrdU pulse-chase experiment in vivo. Analysis of cell counts indicates that the vast majority of memory cells are generated at later time points from cells that have extensively divided.

## Introduction

The number of naive CD8 T cells that are specific for a given pathogen is relatively low, ranging from 100 to 1000 cells (Obar et al., 2008; Haluszczak et al., 2009). Upon infection, these pathogen specific CD8 T cells will be recruited and activated. This, under appropriate conditions, leads to their extensive proliferation and differentiation in a large (10^6^-10^7^ cells) population of effector CD8 T cells that display the capacity to eliminate infected cells. The majority of effector cells will die by apoptosis, except for a smaller subset of memory precursor (MP) cells that will further differentiate to give rise to a long-lived population of memory cells (10^5^ to 10^6^ cells) that will provide protection upon subsequent infection (Murali-Krishna et al., 1998; Crauste et al., 2017). Although these cells are mainly quiescent, they retain the capacity, upon re-exposure to pathogens, to expand and rapidly display effector functions due to epigenetic modifications of genes involved in these processes (Fitzpatrick et al., 1999; Marcais et al., 2006).

In order to better understand the properties of memory cells generated in different settings (Appay et al., 2002), many studies have focused on defining CD8 T cell subsets, relying on a restricted number of surface proteins (Sallusto et al., 1999; Hikono et al., 2007; Jameson and Masopust, 2009). These cell subsets include central and effector memory cells, exhausted memory cells or tissue resident memory cells. Over the years, the study of these subsets has brought a wealth of knowledge on the responsiveness (Wherry et al., 2007; Hikono et al., 2007; Sallusto et al., 1999), homing (Masopust et al., 2001), and self-renewal capacities (Graef et al., 2014; Gattinoni et al., 2012) of memory cells. The molecular pathways sustaining their development have also been largely uncovered. Indeed, the involvement of numerous transcription factors (Intlekofer et al., 2005; Omilusik et al., 2015; Mann et al., 2019; Kaech and Cui, 2012) and epigenetic reprogramming factors (Pace et al., 2018) in the differentiation of different classes of effector and/or memory cells has been described.

The lineage relationship between the different subsets of CD8 T cells (Wherry et al., 2003) and the stage at which activated CD8 T cells diverge from the effector fate to commit to the memory lineage have been extensively studied, with many different experimental approaches leading to results supporting alternative models (Kaech and Cui, 2012). A linear pathway where memory cells are derived from effector cells is supported by early studies using genetic marking of memory cells (Jacob and Baltimore, 1999). A linear model where activated naive cells first differentiate into MP cells that give rise to effector cells has been suggested following in vivo fate mapping of single cells (Buchholz et al., 2013). These early MP cells could correspond to the memory stem cells described in a restricted number of experimental systems (Gattinoni et al., 2012). Fate mapping experiments have highlighted the heterogeneity of effector cells in terms of their functional capacities and their differentiation potential into memory cells (Joshi et al., 2007; Wherry et al., 2007; Sarkar et al., 2008; Kalia et al., 2010). Hence, a new classification of effector cells based on the expression of KLRG1 and CD127 has emerged with, on one side, short-lived effector cells doomed to die at the end of the primary response and, on the other side, MP effector cells that maintain the capacity to differentiate into memory cells (Joshi et al., 2007). In this model and in the first linear models, memory cells are derived from cells that express fully developed effector functions and that have maintained the potential to differentiate into memory cells (Pace et al., 2018; Youngblood et al., 2017). In contrast, a number of other studies have suggested a separation of MP cells at an earlier stage that precedes the differentiation into effector cells. Indeed, branching as early as following the first division has been proposed based on single cell transcriptome analysis (Arsenio et al., 2014; Kakaradov et al., 2017) and would potentially result from an asymmetric division of antigen-triggered cells (Chang et al., 2007). Although these models agree on the early commitment of activated naive CD8 T cells to the memory lineage, there remains some debate about the existence of an early branching (Flossdorf et al., 2015).

More recently, Crauste et al. (2017), based on the modeling of the generation of memory CD8 T cell counts, demonstrated that the total pool of memory CD8 T cells could mainly be generated by a linear pathway; the majority of quiescent memory cells are generated following the transition of naive cells through an early activation effector stage characterized by active cell cycling followed by a late quiescent effector stage (Crauste et al, 2017). In this model, an early branching of memory cells was permitted but it could not account for the generation of the full supply of memory cells.

Overall, functional studies of memory differentiation routes by genetic ablation or cell fate mapping studies have led to the description of multiple possible pathways that lead to a diversity of effector/memory populations. They suggest that memory commitment could take place at several stages of the primary immune response. However, some of these pathways might represent routes followed by only a few cells that generate a minor fraction of the memory cell pool.

In order to uncover the different trajectories followed by naive CD8 T cells to differentiate into memory cells, we have used new trajectory analysis tools that take into account the large amount of information that is delivered by single cell transcriptomics. Indeed, over the last decades, single-cell RNA sequencing (scRNA-seq) has emerged as a powerful tool and allowed a great advance in the exploration of the heterogeneity of the immune system (Papalexi and Satija, 2018). We analysed gene expression dynamics in CD8 T cells collected during the effector response to an acute infection with the Lymphocytic Choriomeningitis Armstrong virus (LCMV-Arm), generated by (Yao et al., 2019) and (Kurd et al., 2020). We applied trajectory inference on these datasets to identify trajectories leading to the generation of MP cells. Using cell-cycle classification and RNA velocity algorithms we show that the differentiation is driven by cell cycle and immune function genes. To identify the molecular regulatory mechanisms supporting the process, we then performed a gene regulatory network (GRN) inference analysis which allowed us to identify a cluster enriched in cells harbouring transcription factors associated with MP cells. Using a MP gene signature, we confirmed that this cluster was enriched in MP cells, though cells expressing that signature were also found at multiple points along the trajectory. Finally, we used another pathogen infection and BrdU labelling to generalise and validate these results in vivo. Analysis of cell counts confirmed that although memory cells are generated continuously all along the trajectory, the majority of memory cells were derived from cells that had proliferated and acquired effector functions.

## Results

### Trajectory inference of the CD8 T cell response to an acute infection

In order to gain insight into the differentiation dynamics of CD8 T cells in response to an acute infection (LCMV-Arm), we performed trajectory inference on a scRNAseq dataset generated by Yao et al (2019) using two recently published methods, Slingshot (Street et al., 2018) and TinGa (Todorov et al., 2020). This dataset consists of measurements on 20,295 splenic CD8 T cells generated following LCMV-Arm acute infection and isolated at two different time points (4.5 and 7 days post infection (dpi)), in two separate replicates. We identified the 2,000 most highly variable genes in the dataset using variance modelling statistics from the Scran R package (Lun et al., 2016), on which we applied both trajectory inference methods. Slingshot was shown to be very efficient in a study that compared more than 40 methods on a large number of datasets (Saelens et al., 2019). TinGa is a new method for trajectory inference that showed comparable results on simple trajectories and better results on complex trajectories than Slingshot (Todorov et al., 2020). These two methods both share a first step in which the dimensions of the data are reduced, either by principal component analysis for Slingshot, or by multidimensional scaling (MDS) for TinGa. In the two resulting representations of the data, the cells form a continuum from cells taken 4.5 dpi to cells taken 7 dpi (Figure 1A and Supplementary figure 2A).

**Figure 1:**
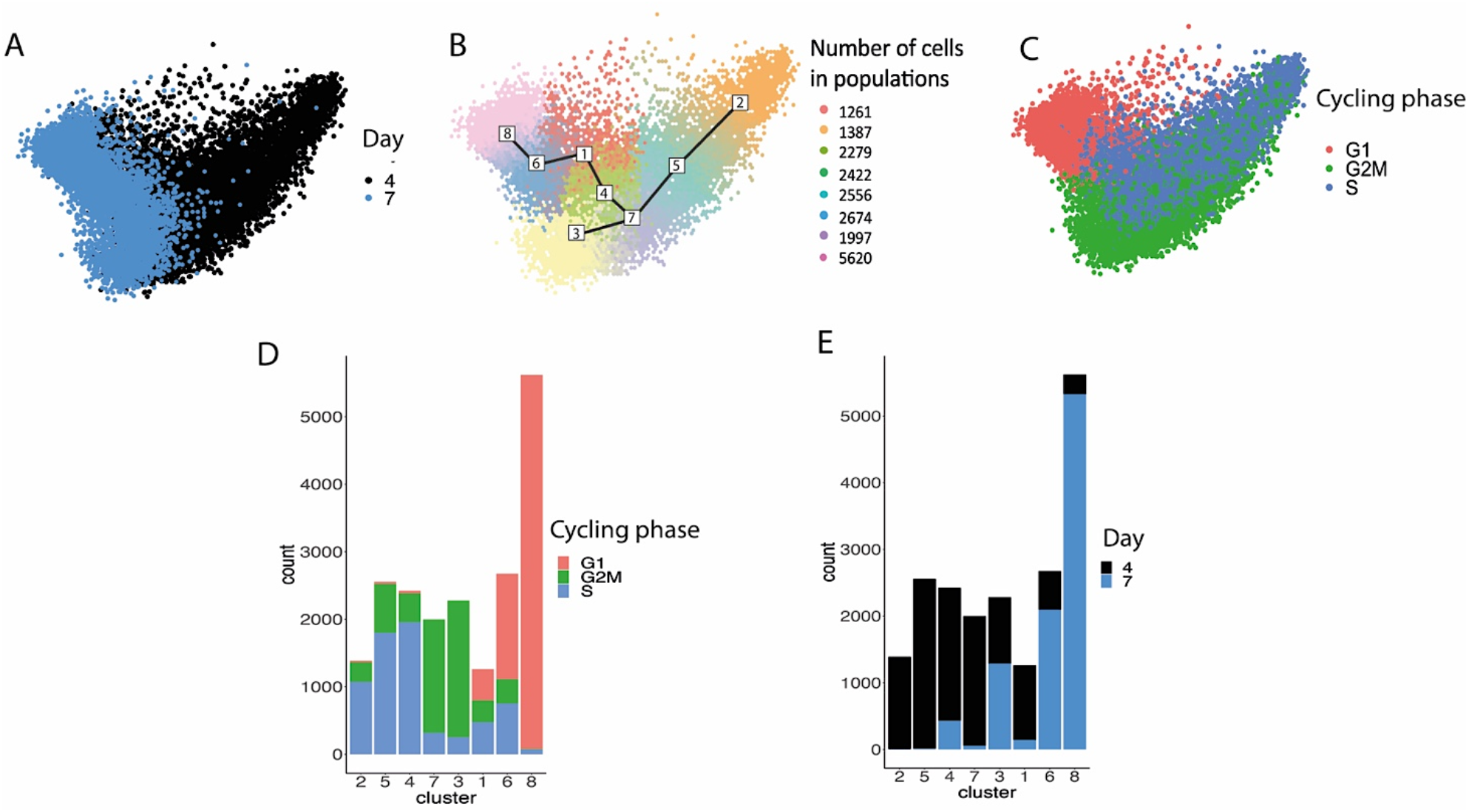
TinGa trajectory inference **(A-C)** Visualization of the cells in a 2D space computed by multi-dimensional scaling **(A)** The cells were colored according to the two experimental time points 4.5 and 7 days post LCMV-Armstrong infection. **(B)** The TinGa algorithm identifies a branching trajectory in the data, that is represented by a black line. Milestones along the trajectory can be used to define subgroups of cells that are represented by different colors. They will be referred to as “clusters”. The number of cells in each cluster are indicated. **(C)** The cells were classified into one of the cycling phases (either G1, S, or G2/M) using the Seurat package and colored accordingly. **(D-E)** The number of cells in the G1, S and G2/M phases **(D)** and in the two experimental time points **(E)** are shown for each cluster.

Slingshot first applies clustering to the data and then identifies transitions between these clusters. It identified a linear trajectory starting among cells from day 4.5 post-infection (pi), transitioning through a mix of cells from day 4.5 to 7 pi, and ending in a part of the data that was enriched in cells from day 7 pi (Supplementary figure 2B). The genes that varied the most along this trajectory are identified in Supplementary figure 2C. The linear Slingshot trajectory seemed to start in early activated cells, in which we identified an overexpression of Ybx1, Rps2, Rps8 genes involved in the initiation of transcription. The trajectory then transitioned through a state where the cells seemed to be undergoing divisions (Tubb4b, Tuba1b, Ccna2, Cks1B genes) and ended in cells that expressed genes associated with immune functions (such as Ccl5, Hcst, B2m, H2-D1). In comparison, the trajectory identified by TinGa started similarly to the Slingshot trajectory, but then split into two branches (Figure 1B). One small branch (identified by the number 3) corresponded to cells that seemed to be in a highly cycling state, whereas the other longer branch ended, after several transitional states, in the effector-memory-like state described in the Slingshot trajectory (Supplementary figure 1A). Eight transitional states were identified along the TinGa trajectory. For convenience, these eight transitional populations will be referred to as clusters from now on.

Among the 40 genes that varied the most along the two trajectories defined by Slingshot and TinGa (Supplementary figure 1 and 2C), 33 were commonly found in both trajectories. This suggests that, even though TinGa identified an extra small branch that Slingshot included in a linear trajectory, the genes that are mainly driving cells along the two trajectories are similar. Interestingly, when we applied Slingshot and TinGa to a reduced set of 1,300 highly variable genes, both methods recovered a branching trajectory (Supplementary figure 1B and Supplementary figure 2D). This indicates that the main trajectory uncovered is robust and that the small-branch identified differs only marginally from the neighboring cluster.

### The inferred trajectories retrace an early-late-memory differentiation pathway

To further characterize the CD8 differentiation trajectory supported by the single cell transcriptomics data, we used the trajectory obtained with TinGa as it identified more transition points along the route and hence might give a more refined definition of differentiation steps. As both the Slingshot and the TinGa trajectories were clearly driven by cell cycle associated genes (Supplementary figure 1A and Supplementary figure 2C), we used a classifier from the Seurat R package (Tirosh et al., 2016) to allocate cells to the G1, S or G2/M cell-cycle phases (Figure 1C). We identified clear cycling trends along the trajectory. Cells in clusters 2, 5 and 4 were mainly classified in the S phase (Figure 1C-D) while clusters 7 and 3 were de facto classified in the G2/M phase (Figure 1C-D). Cluster 6 was clearly enriched in G1 cells, while cluster 8 contained almost exclusively cells in G1 (Figure 1C-D). Hence, these results showed that the Tinga trajectory consisted of a first cycling effector population that differentiated in a quiescent effector population. Interestingly, TinGa identified three clusters enriched in cells positioned in the S phase (cluster 2, 5 and 4) and two clusters enriched in cells positioned in the G2/M phase (cluster 7 and 3). These clusters, however, differed in terms of sampling days and the two clusters (3 and 4) positioned at a later pseudotime by TinGa contained a larger fraction of cells sampled on day 7 compared to the earlier clusters with a similar cell cycle position. Thus, to unravel genes driving the trajectory, while overcoming cell cycle gene expression biases, we performed a differential expression analysis between cells from the same cycle phase present in each neighboring cluster along the trajectory (Supplementary Table 1 and Supplementary figure 3A). This highlighted the slow transition from early activation markers (Xcl1, Srm) to effector functions (Ccl5, Id2, Gzma/k) and quiescence (Btg1) (Supplementary figure 3A right panel).

To further define the dynamics of cell differentiation, we applied the scVelo algorithm (Bergen et al., 2020) that defines RNA velocities. These were projected onto the TinGa embedding (Figure 2A). ScVelo retraced two RNA trajectory dynamics (Figure 2B). The first suggests a circular trajectory that would fit with cells going through the cell cycle. The second corresponds to a linear trajectory of differentiation leading from clusters 5 and 7 to 8. Interestingly, the cells of cluster 3 seemed to join those of cluster 8. Thus, the small branch identified by TinGa could correspond to a transient state composed of cells passing from a state of proliferation to a quiescent state.

**Figure 2:**
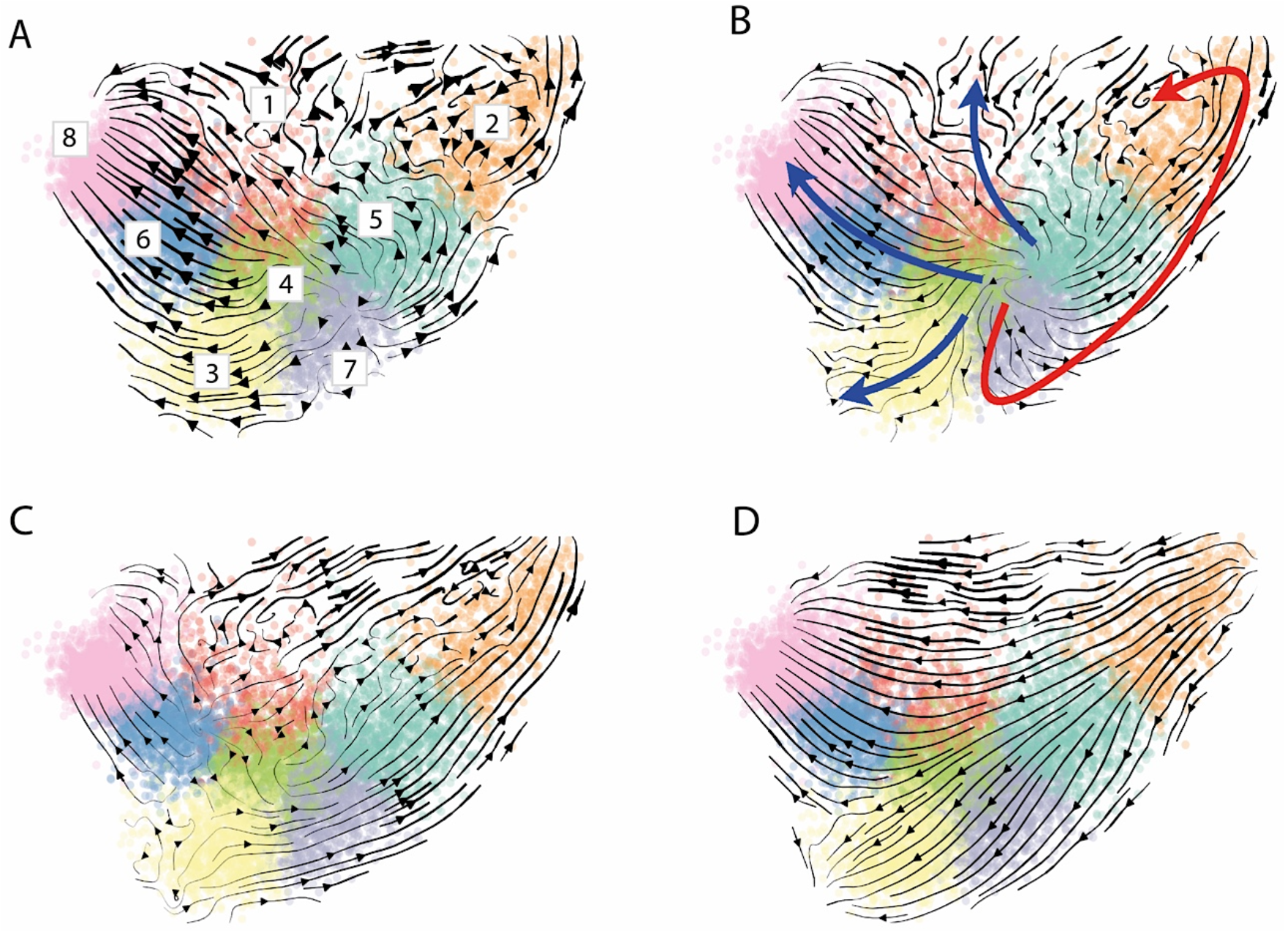
Gene expression dynamics along the differentiation trajectories RNA velocities are projected onto the TinGa embedding. The cells in the trajectory were colored according to their TinGa milestones. **(A)** Velocities were calculated using all genes. Numbers correspond to TinGa clusters. **(B)** The RNA velocities show two distinct dynamics. In clusters 2, 5 and 7, cells are cycling (red arrow) but can commit to the differentiation dynamic (blue arrows) by leaving clusters 5 and 7 to reach cluster 8. **(C-D)** RNA velocities based only on cell cycle and migration **(C)** or immune function-related **(D)** genes.

A similar dynamic was obtained using only the top 50 genes contributing to the scVelo’s dynamical model (Supplementary figure 3B, Supplementary Table 2) indicating that they were sufficient to recover the overall cell dynamics (Figure 2A). We then analyzed the molecular function associated with these 50 genes and found that they could be broadly classified into three categories (cell cycle, migration and immune function) (Supplementary Table 2). To compare the contribution of these genes to the dynamic, the RNA velocities associated with cell cycle/migration related genes or immune function genes were calculated and projected onto the TinGa embedding (Figure 2C-D). The cell cycle and migration genes clearly defined the first circular dynamics found at the start of the trajectory and also contributed to the differentiation process (Figure 2C) while the immune genes recapitulated a linear trajectory going from cluster 2 to 8 (Figure 2D). By looking at individual gene dynamics, we found that genes act on different parts of the cell differentiation trajectory. Genes such as Id2 have an early effect, with stronger contribution to the global velocities in clusters 4, 5 and 7, while genes such as IL18r1 and Gimap6 start to act at later pseudotime with stronger velocities in the final clusters of the trajectory (Supplementary figure 3C).

The trajectory inference based on single cell transcriptomic data seems to recapitulate the two effector compartments that we have previously described, i.e., a first set of early effector cells that are cycling followed by a set of late effector cells that are quiescent and express increased levels of genes encoding immune effector functions (Crauste et al., 2017).

### Gene regulatory interaction analysis

To further characterize the transitional stages defined along the TinGa trajectory, we identified regulatory interactions between transcription regulators and their target genes in the dataset using the BRED tool (Cannoodt et al., 2019). We identified six main GRN-modules, that we defined as groups of target genes gathered around a regulator (Figure 3A). As expected, based on previous results on the cell cycle, three of these modules (Pcna, Hmgb2, Cenpf) were strongly enriched in genes involved in cell cycle regulation. The Ybx1 GRN-module contained two groups of genes. One coding for proteins involved in RNA and protein synthesis metabolism that were up-regulated in the cells from the cluster 2-5-7 branch, the other for immune functions that were enriched in clusters 6 and 8 (Supplementary figure 4). Two GRN-modules were composed essentially of genes associated with the immune response. The GRN module Spi1 was expressed in very few cells along the trajectory (Supplementary figure 5). In contrast, the Tcf7/Id2/Phb2 GRN-module contained genes coding for transcription factors and immune functions, associated with the differentiation of CD8 T cells in memory cells. These genes were expressed in different clusters along the trajectory (Supplementary figure 6). Interestingly, cluster 1 was enriched in cells that coexpressed genes from the Tcf7/Id2/Phb2 modules which were associated with a MP cell phenotype as defined by a number of studies (Yao et al., 2019; Wu et al., 2016; Utzschneider et al., 2016; Chen et al., 2019). Indeed, they expressed Tcf7 and Id3, two transcription factors that were previously associated with a MP potential (Yao et al., 2019). Two target genes, Slamf6 and Tnfsf8, were found to be positively correlated with the presence of Tcf7 in the Tcf7/Id2/Phb2 module. In contrast, the Id2 transcription factor, that has previously been associated with an effector fate (Omilusik et al., 2018), was repressed in these cells, as was the effector associated gene Gzmb (Figure 3B and Supplementary figure 6).

**Figure 3:**
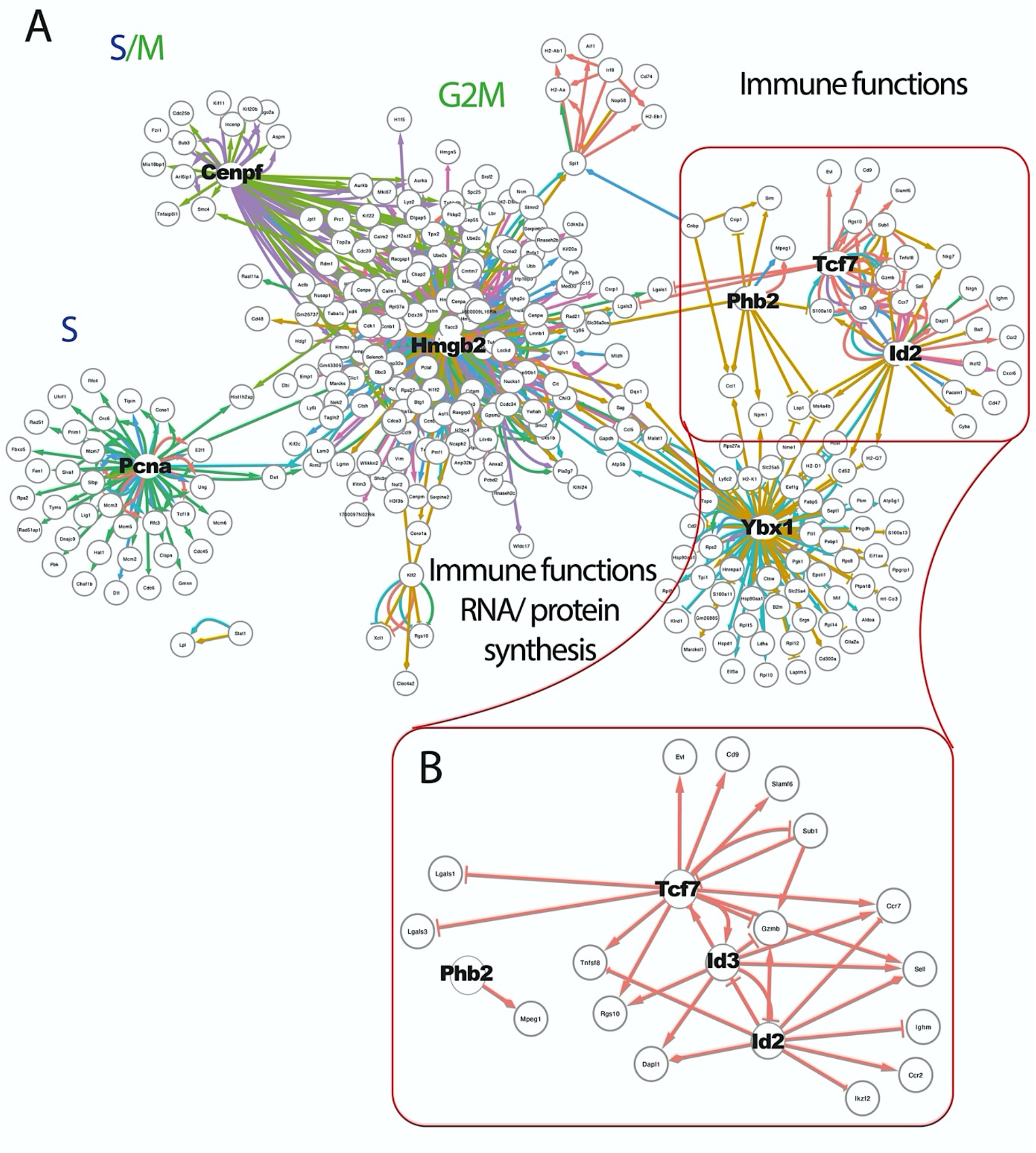
Gene regulatory interactions **(A)** Gene regulatory network identified with BRED. In this GRN, regulatory processes are symbolized by arrows that are directed from transcription factors to their target genes, or to other transcription factors. The top 100 interactions per TinGa cluster are represented, and are colored according to the cluster in which they are occurring. The shape of the arrow indicated whether the interaction was an activation (->) or an inhibition (-|). **(B)** Zoom on the Id2/Tcf7/Id3 module identified by BRED. Only the interactions occurring in cluster 1 in the TinGa trajectory are represented.

In summary, cluster 1 seemed to contain an interesting set of cells in which effector functions were down-regulated, while genes associated with a memory precursor signature were overexpressed. We thus decided to further characterize the cells in cluster 1.

### TinGa identifies distinct clusters associated with a memory-precursor phenotype

Cluster 1 was mainly composed of cells from day 4.5 pi, a large fraction of which (40%) was classified as being in the G1 phase of the cell cycle (Figure 1D-E, Supplementary figure 7A). This contrasted with other clusters enriched in cells from day 4.5 pi, such as clusters 2 and 5, that contained very few cells classified as being in G1 (Figure 1D-E).

To ascertain that cells in cluster 1 had been activated, we compared their transcriptome with the genes expressed in cluster 2 positioned at the beginning of the trajectory. Results in Supplementary figure 7B showed that the cells in cluster 1 expressed an increased amount of genes coding for effector functions such as Ccl5 and Gzma compared to cluster 2, indicating that these cells had been activated as they had started to acquire effector functions. Cells in cluster 1 also expressed interferon-induced genes such as Ifl27ia, Ifl203, Ifl47 (colored in red in Supplementary Fig 7B), indicating that these cells had responded to the pathogen-induced innate response. We thus concluded that cluster 1 contained cells from day 4.5 pi that had been activated but already displayed traits of quiescence.

Cluster 1 cells expressed increased amounts of Tcf7, Id3 and Ltb as compared to all other cells in the trajectory, while Klrg1, a gene associated with terminal differentiation, was down-regulated in these cells (Supplementary figure 7C). This was in agreement with the activation of the Tcf7/Id2/Phb2 module, containing genes associated with a MP potential in these cells. To confirm the MP genetic program of cells in cluster 1 and to identify all MP cells along the trajectory, we performed a gene set enrichment analysis (GSEA) using the MP gene signature recently defined in (Yao et al., 2019) (Supplementary Table 3 and Figure 4A). We identified 833 MP cells that were mainly localized in clusters 1 and 8 (Figure 4A-B). Unsurprisingly, cluster 1 was the most enriched in the MP signature with 15% of the cells presenting the signature. Cluster 8 also contained a significant fraction (9%) of MP cells. However, the majority of MP cells were associated with cluster 8 that contained 3 times more MP cells than cluster 1 (Figure 4B-C). The majority of MP in cluster 1 and 8 were associated with the G1 phase of the cell cycle compared to those in the other clusters that were mainly in S and G2/M phase (Figure 4C).

**Figure 4:**
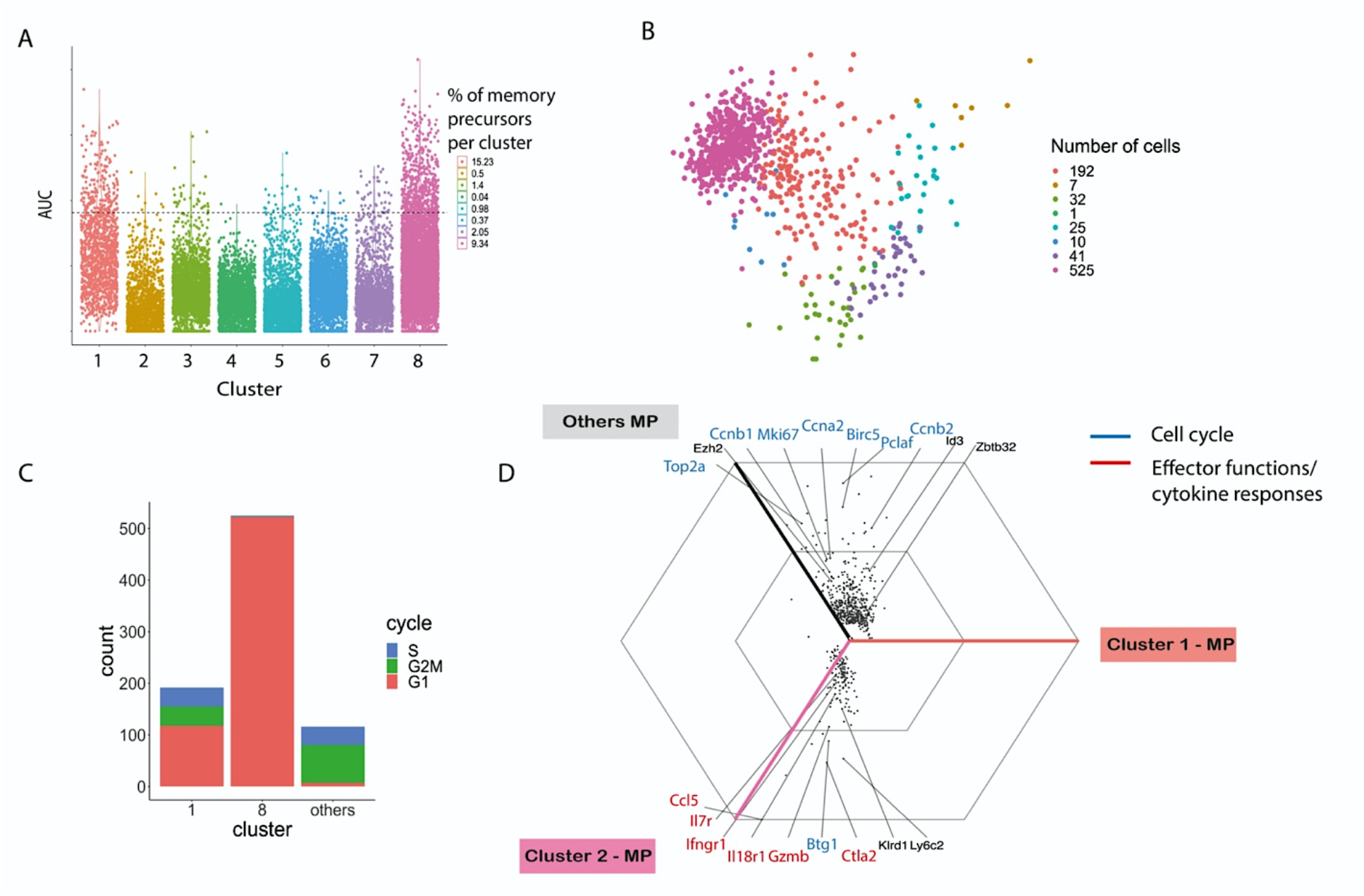
Memory precursor cell identification and characterization. **(A)** Memory precursor signature enrichment in each cluster along TinGa’s trajectory. The cells above the threshold represented as a dotted line are considered as memory precursors. The percentage of cells with a memory precursor signature in each cluster is indicated. AUC: area under the curve **(B)** The cells with a memory precursor signature were represented on the TinGa map and colored according to the cluster they came from. The number of cells with a memory precursor signature in each cluster is indicated. **(C)** Distribution in the G1, S and G2/M cell cycle phases of cells with a memory precursor signature from clusters 1, 8 or all others. **(D)** Triwise plot of the log fold-change expression of genes that were differentially expressed between the memory precursor cells found in cluster 1, 8, and all other memory precursors. Only the genes that were differentially expressed with a p-value < 0.05 are represented. The internal hexagon corresponds to a log fold-change of 1, the external hexagon corresponds to a log fold-change of 2.

To determine the number of MP cells that had effectively been found on each sampling day, we recalculated the number of MP cells present in the spleen of animals at the two experimental time points (see Methods section). Based on the number of LCMV-Arm specific CD8 T cells present in the spleen on day 4.5 and 7 pi, we could estimate the number of cells with a MP gene signature on these two days to be 3,850 and 643,000. This indicated that the number of MP cells generated 4.5 days after infection is more than 150 times lower than the number of MP cells generated 7 days after infection, in agreement with values estimated in (Crauste et al., 2017).

To investigate differences between MP cells generated at day 4.5 and at day 7 pi, we compared the gene expression profiles of cluster 1 to cluster 8 MP cells respectively, and to the profiles of MP from all the other clusters (Figure 4D). Both clusters 1 and 8 MP cells showed a decreased expression of genes driving the cell cycle compared to the other MP, in agreement with their position in G1 phase (Figure 4C-D). Cluster 8 MP cells have an increased expression of genes coding for proteins involved in effector functions (Gzmb, Ctla2, Ccl5) or cytokine response (Il7r, Il18r1, Ifngr1) compared to cluster 1 MP cells, indicating that, although they display a MP gene signature, they have also acquired effector cell properties. This was in agreement with the data showing that effector cells could dedifferentiate into quiescent memory cells (Youngblood2017). Interestingly, cycling MP (i.e., MP from clusters other than 1 and 8) expressed genes coding for transcription factors (Zbtb32, Id3) or histone modifier (Ezh2) involved in the regulation of the developmental switch between effector and MP cells, suggesting that cycling MP are still oscillating between these two fates (Kakaradov et al., 2017; Shin et al., 2017; Yang et al., 2011).

To confirm the continuous generation of MP cells, we analyzed a second transcriptomic dataset generated by Kurd et al. that contained cells sampled at multiple time points during the primary response (day 4, 5, 6, 7 and 10 post-LCMV infection). Highly variable genes expressed by the 9,614 cells were selected and TinGa was applied (Supplementary figure 8A). The trajectory obtained is similar to the Yao et al.’s data (Yao et al., 2019) with the first part of the trajectory being enriched in cycling cells (cluster 1,3, 8, 6 and 4) which were sampled on day 4, 5, 6 and 7 pi. The second part contained quiescent cells sampled on day 6, 7 and 10 pi (Supplementary figure 8A-B-C). Similarly, 574 MP cells were found by GSEA all along the TinGa trajectory (Supplementary figure 8D-E).

Overall, these results suggested that MP cells with different functional and differentiation statuses, from activated cycling cells to quiescent effector cells, were present at different points along the trajectory.

### In vivo validation of memory cell generation at different time points following activation of CD8 T cells

Our in-silico analyses strongly suggest that CD8 T cells become quiescent and differentiate into memory cells at different stages following activation in response to acute viral infection. To validate this result in vivo, we used BrdU pulse-chase experiments. Indeed, these allow tracking cells that proliferate during the pulse time and stop soon thereafter, thus maintaining their BrdU labelling in the memory phase. This way we can identify MP cells present at the time of pulse. We also wanted to extend the data to other experimental systems so we used vaccinia virus (VV) that induces a local acute infection in the lung when inoculated intranasally. Thus, mice were infected intranasally with a VV harboring the NP68 epitope and we followed the activation of TCR transgenic F5 cells (Crauste et al., 2017). Mice were given one injection of BrdU on days 4, 7 and 11 pi and BrdU labelling was analyzed after 24 hours in the spleen and the lymph nodes draining the lung and nasal cavity (Supplementary figure 9A).

Following VV infection, TCR transgenic F5 CD8 T cells increased in proportion and numbers over time in both spleen and draining lymph nodes (dLN), with a peak 8 dpi (Figure 5A-B). The percentage of BrdU labelled cells, representative of proliferating CD8 T cells, was maximal 5 dpi in the dLN and 8 dpi in the spleen, reflecting the local initiation of the CD8 T cells immune response following intranasal infection (Figure 5C-D). The number of BrdU labelled cells was maximal both in spleen and dLN 8 dpi and started to decrease thereafter with a limited number of cycling cells detected 12 dpi.

**Figure 5:**
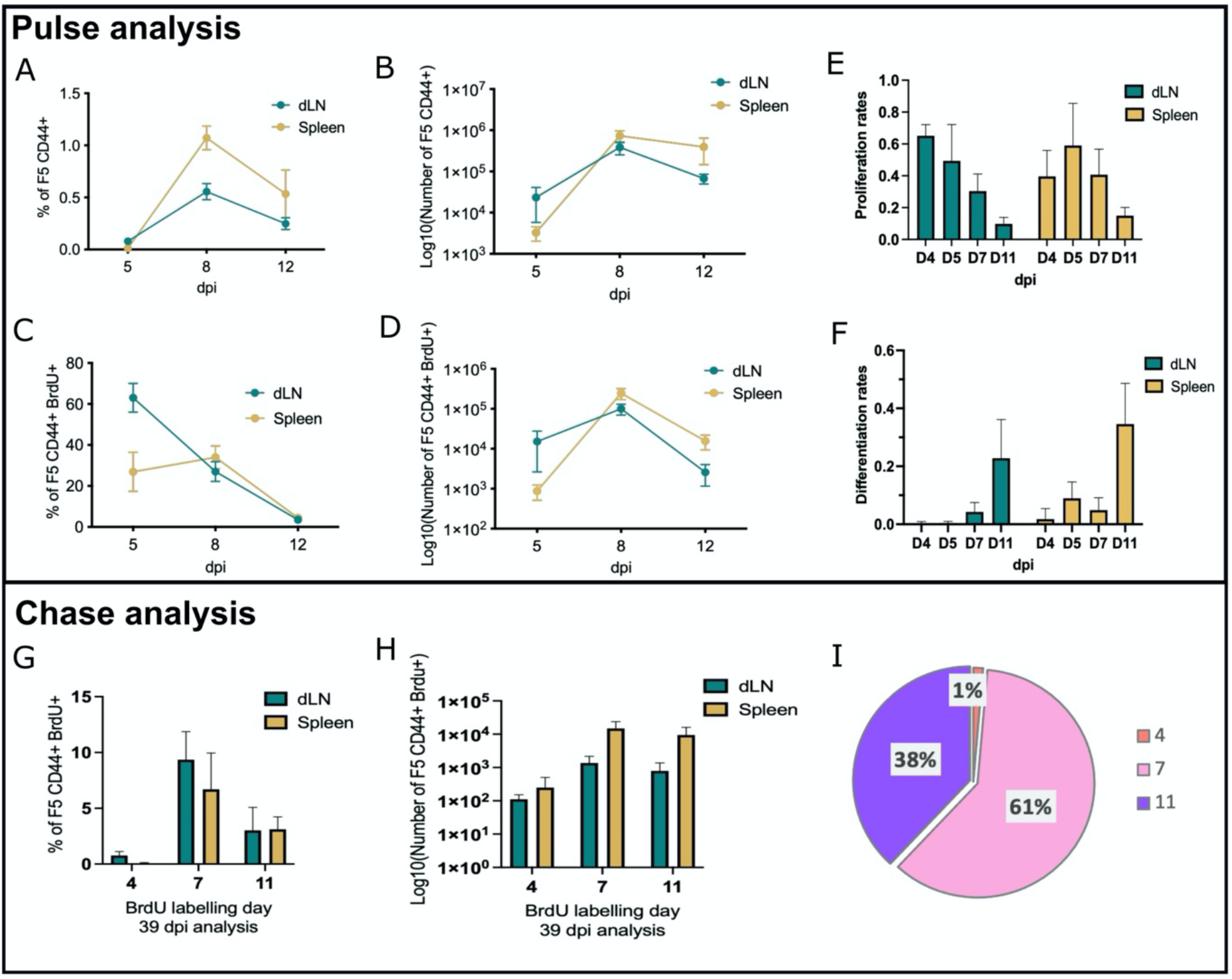
In vivo identification of memory precursors using BrdU labelling. **(A-D)** The percentages **(A, C)** and numbers **(B, D)** of total F5 CD44+ **(A, B)** and BrdU+ F5 CD44+ **(C, D)** CD8 T cells were determined 24h after BrdU injection (pulse) in the indicated organs. **(E)** Proliferation rates of Early Effector (EE) cells in the indicated organs from day 4 to day 11 post-infection. **(F)** Differentiation rates of EE cells into Late Effector (LE) cells in the indicated organs from day 4 to day 11 post-infection. **(G-H)** The percentages (**G**) and numbers (**H**) of BrdU+ F5 CD8 memory T cells originating from cells labelled on days 4, 7 or 11 pi was determined 39 days after BrdU injection (chase) in the indicated organs. **(I)** Proportion of memory cells originating from MP labelled at days 4, 7 or 11 pi normalized to the total number of recovered BrdU* F5 CD8 memory T cells. Data are representative of 3 independent experiments.

Using data from three independent experiments we next estimated (see Methods section) the proliferation and differentiation rates of cycling effector cells (Crauste et al., 2017) on days 4, 5, 7 and 11 pi. In agreement with the BrdU labelling profile of total CD8 T cells we found that the proliferation rate of effector cells peaks on days 4-5 before quickly decreasing both in dLN and spleen (Figure 5E). This is in agreement with previous results (Crauste et al., 2017) obtained on blood samples. In contrast, their differentiation rates to quiescent effector cells are very low on days 4-5 pi, increasing on day 7 pi with the highest rate being observed on day 11 pi, once proliferation has drastically vanished, in both spleen and dLN (Fig 5F).

We then measured the fraction and number of BrdU-labelled CD8 T cells in the memory phase (39 dpi) in order to evaluate the MP cells present on day 4, 7 or 11 pi of the response (Figure 5G-H). As predicted by the single cell transcriptomic data, we found that memory cells could derive from activated/effector cells at all investigated times. However, the largest fraction of memory cells was derived from cells labelled on day 7 or later (Figure 5I). Importantly the few cells that were labelled with BrdU between days 11 and 12 gave rise to a significant fraction of the memory cell pool in agreement with their increased differentiation rates (Figure 5F).

Finally, we compared protein expression of memory cells generated on day 4.5 or day 7 pi. We thus performed a BrdU chase experiment (Supplementary figure 9B) and measured the expression of proteins encoded by genes that were differentially expressed in the single cell transcriptomic dataset (Figure 4D and Supplementary Table 4). We found that CCL5, which was the most differentially expressed gene in late d7 MP cluster 8 (Figure 4D), was also expressed at a higher protein level by F5 memory cells generated at 7 dpi (Figure 6A). The expression of CCL5 was also measured on endogenous antigen-induced BrdU positive memory cells, identified based on their CD49d expression (Grau et al., 2018) (Supplementary figure 9C). Similarly, we found a significant increase of CCL5 expression on BrdU+ endogenous memory cells generated at 7 dpi in the dLN and spleen (Figure 6B-C).

**Figure 6:**
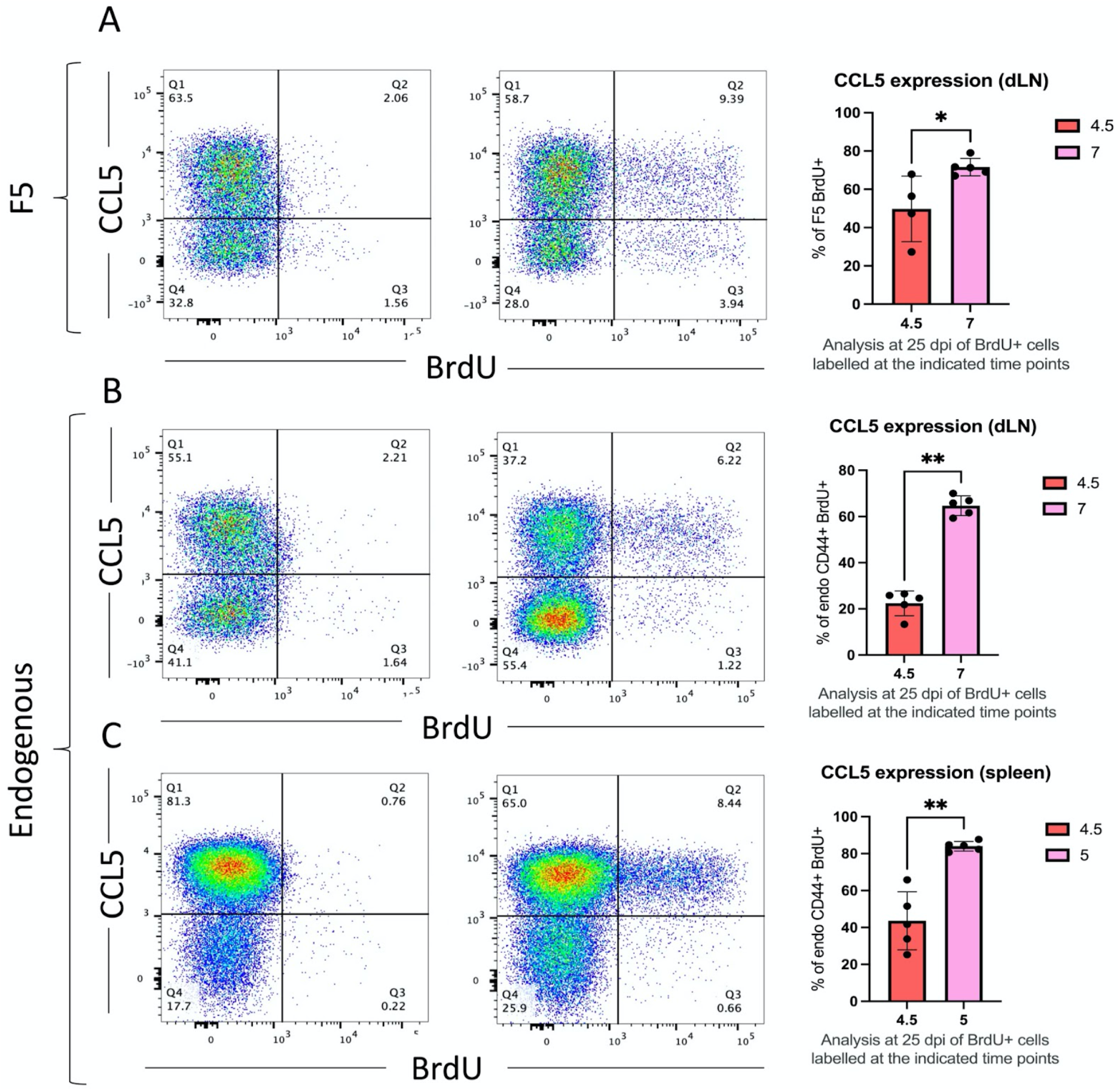
CCL5 expression of early and late generated memory cells by flow cytometry. **(A)** Flowcytometry plots and quantification of BrdU+ F5 memory cells expressing CCL5 labelled on day 4.5 and 7 with BrdU was assessed. **(B, C)** Flow-cytometry plots and quantification of CD44+ CD49d+ BrdU+ endogenous memory cells expressing CCL5 labelled on day 4.5 and 7 with BrdU was assessed in the draining lymph node (**B**) and in the spleen (**C**). *p < 0.05 (Mann-Whitney test). Data are representative of 2 independent experiments.

Overall, these results show that although memory cells are generated continuously after activation, the majority of memory cells are generated late during the effector phase (Figure 7).

**Figure 7:**
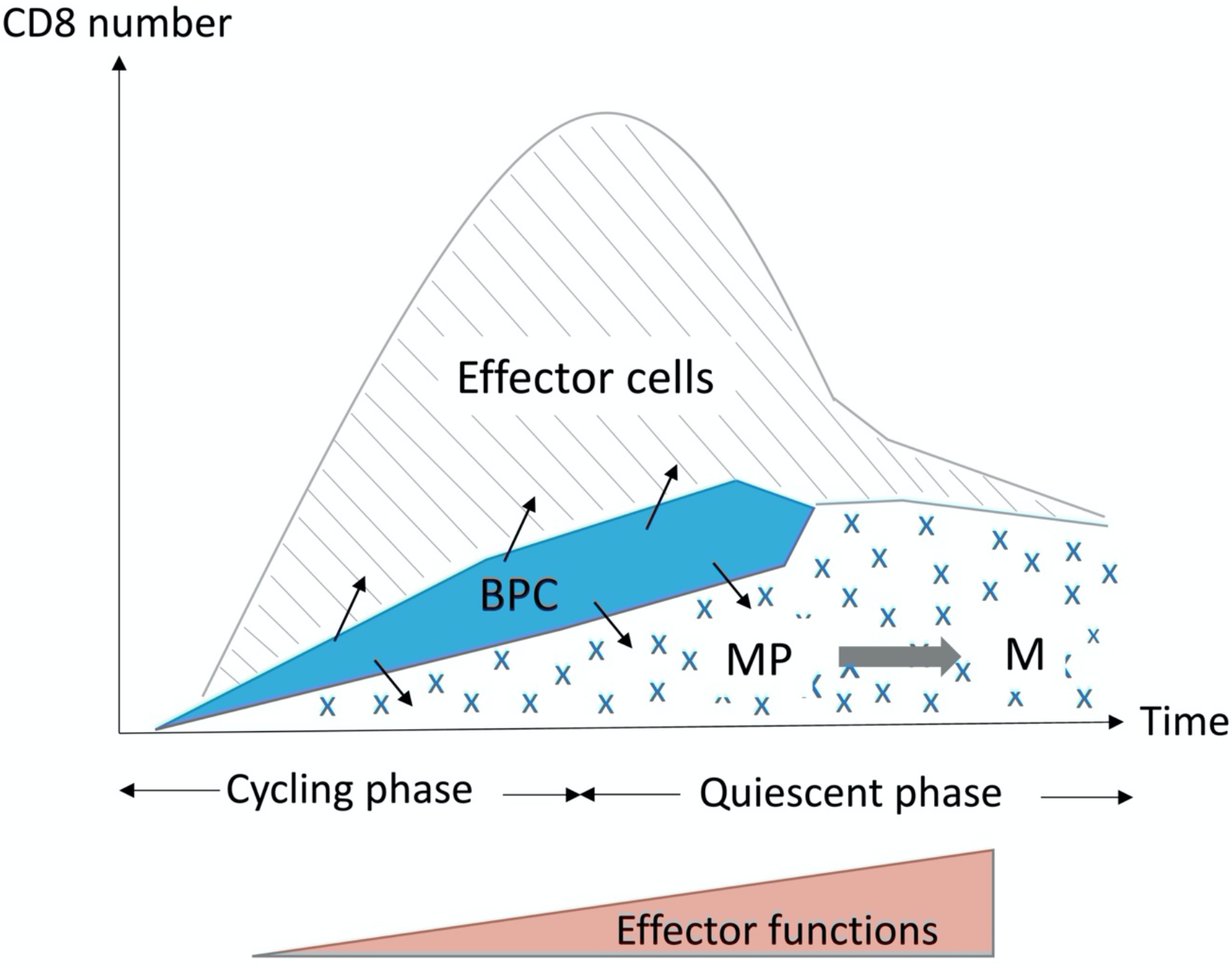
Using in silico analysis of single cell transcriptomics and in vivo tracing of memory precursors coupled to mathematical modeling, we demonstrate that MP are generated continuously during the primary response with the largest fraction being generated at the peak of the expansion phase. BPC: bipotent precursor cells; MP: memory precursors; M: memory

## Discussion

In this study, we have used trajectory inference tools to analyze the generation of memory precursor CD8 T cells during a primary response against an acute viral infection. A single cell transcriptomic dataset (Yao et al., 2019) generated at two timepoints during the primary response, was analyzed using two recently developed trajectory inference algorithms (Slingshot (Street et al., 2018) and TinGa (Todorov et al., 2020)). These tools allow modeling gradual transitions between cell states, as they tend to preserve the local similarities between cells, thus predicting the likely differentiation path followed by cells activated *in vivo* by the virus (Saelens et al., 2019). Trajectory inference tools have become essential as they allow to predict the fate of cells that have to be lysed to analyze their cellular content and/or transcriptome. Although different dimensionality reduction and trajectory computation approaches were used, the trajectories identified by both algorithms were driven by similar sets of genes and displayed a consistent trajectory starting among cells from day 4.5 postinfection that were mainly cycling and ending among cells from day 7 post-infection that were mainly quiescent. Importantly, there was a significant overlap between cells collected on each day as clusters in the middle of the trajectory contained cells from both time points. This indicates that the differentiation process although continuous is heterogeneous in its duration as for example some cells exit the cell cycle at early time points or acquire effector functions more rapidly. This is in agreement with experiments tracking the fate of single T cells in mice that have shown that the clonal size of memory cells generated from a naïve CD8+ T cell is heterogeneous (Buchholz et al., 2013; Gerlach et al., 2013). The trajectory identified by TinGa was more refined as it identified 8 transitional stages, one of which (cluster 1) was strongly enriched in MP cells identified using a gene signature derived from Yao et al. (2019).

We also applied scVelo (Bergen et al., 2020), a method that uses the splicing state of transcripts to calculate RNA velocities. The projection of RNA velocities on the TinGa-generated map evidenced two cellular behaviors with early cycling cells that remain on a circular trajectory and later cells that follow a linear path. These two behaviors were associated with cell cycle and immune function genes, respectively. Importantly, the linear trajectory driven by the immune effector genes started in early (d4.5) cells underpinning cycling and quiescent cells, thus reflecting the progressive expression of effector functions by activated CD8 T cells. These results are in agreement with the two effector compartments previously described, namely the early cycling effector cells and late quiescent effector cells expressing genes encoding immune effector functions, through which most MP cells have to go to generate the full pool of memory CD8 T cells (Crauste et al., 2017).

We herein found that MP are present at all pseudo-times, with an enrichment in clusters 1 and 8. The majority of MP cells in clusters 1 and 8 were in the G1 phase of the cell cycle suggesting that they were on their way to become quiescent memory cells. We confirmed the continuous generation of MP cells on another dataset from Kurd, et al. (2020). Importantly, we estimate that the number of MP cells generated on day 7 pi is around 100-fold higher than the number generated on day 4.5 pi. In vivo pulse-chase BrdU experiment confirmed that CD8 T cells became quiescent memory cells at different stages of an acute infection and that the differentiation rate of early effector cells increased over time. Overall, our data support a model where MP cells are generated continuously over the duration of the expansion phase and beyond, with the majority generated at the peak of the response. Memory precursors identified on day 7 (cluster 8) differ from MP cells generated earlier in the response, mainly by their expression of genes coding for CD8 effector functions (Gzmb, Ccl5) and we confirm that CCL5 is expressed at higher protein level by memory cells generated at 7 dpi compared to cells generated at 4.5 dpi. This is in agreement with the gradual acquisition of epigenetic modifications that lead to a poised transcriptional state of the effector molecule loci in memory CD8 T cells (Dogra et al., 2016; Henning et al., 2018). Based on differential gene expression, we searched for surface markers that could distinguish memory precursor CD8 T cells generated early or late in the response. Unfortunately, we have been unable to identify such markers which would have allowed us to compare the functions and self-renewal capacities of these cells.

The continuous generation of MP over the duration of the effector phase could be explained by the sustained proliferation of MP generated early in the response. These cells would maintain self-renewing capacity while opening the chromatin at effector function gene loci. This would fit with the increased expression of mRNA coding for effector functions in MP identified on day 7. We estimate that cycling MP cells represent only about 15% of all MP cells. Interestingly, these cells differ from quiescent memory precursors by the expression of the transcription factors Zbtb32 and Ezh2, which encodes a catalytic subunit of the polycomb repressive complex 2 (PRC2) (Gray et al., 2017). Zbtb32, which is transiently expressed during the effector phase has recently been shown to control the magnitude of effector cells and the generation of memory cells (Shin et al., 2017). Epigenetic modification by Ezh2 controls the survival and cytokine production of effector cells. Also, it would be involved in the developmental switch between terminal effector cells and memory cells by depositing H3K27me3 in T effector cells (Gray et al., 2017; Kakaradov et al., 2017). Thus, proliferating MP cells could represent bipotential cells that oscillate between two fates: the terminally differentiated effector fate that is associated with the repression of the self-renewing capacity and the activation of effector function loci and the memory precursor fate that maintains the self-renewing capacity while acquiring bivalent chromatin modification marks on gene encoding effector functions. This hypothesis would be in line with a recent study by Pace et al. (2018) suggesting that cycling cells may represent bipotent differentiation intermediates expressing both effector and stem/memory potential. A similar differentiation pattern has recently been found in a hematopoietic stem cell differentiation model (Moussy et al., 2017). Importantly in that model and similarly to our data, the number of divisions performed by bipotent cells before arresting and stabilizing in one or the other fate is heterogenous.

Such a continuous bivalent model could reconcile a number of previously proposed conflicting models that positioned memory precursor cells at either early or late stages following activation (Arsenio al., 2014; Buchholz al., 2013; Flossdorf al., 2015; Jacob and Baltimore, 1999; Kakaradov al., 2017) (Figure 7). Importantly, it could account for the diverse sizes of clones derived from a single cell, observed in fate mapping experiments (Buchholz et al., 2013; Gerlach et al., 2013) while being in agreement with the dynamical modelling of memory CD8 T cells generation (Crauste et al., 2017). Finally, it would allow the deposition of epigenetic fingerprints on genes that encode effector functions and are poised for rapid expression in memory cells.

## Methods

### Experimental procedures

#### Mice

C57BL/6J mice were purchased from the Charles River Laboratories. F5 TCR [B6/J-Tg(CD2-TcraF5,CD2-TcrbF5) 1Kio/Jmar] transgenic mice were provided by Prof. D. Kioussis (National Institute of Medical Research, London, U.K.) and backcrossed on CD45.1 C57BL/6 background (Jubin et al., 2012). Mice were bred or housed under specific pathogen free conditions in our animal facility (AniRA-PBES, Lyon, France). All experiments were approved by our local ethics committee (CECCAPP, Lyon, France) and accreditations have been obtained from governmental agencies.

#### BrdU labelling

Mice received 2.10^5 naive CD45.1 F5-Tg CD8 T cells by intravenous (i.v.) injection one day prior intranasal (i.n.) infection with VV-NP68 (2.10^5 pfu under 20 \muL). Mice then received one intraperitoneal (i.p.) BrdU injection (2 mg, Sigma). BrdU labelling was analyzed 24h after BrdU administration or 25 and 39 days post infection (dpi).

#### Cell analyses

Mice were sacrificed by cervical dislocation and spleen and draining lymph nodes (cervical and mediastinal) were collected. Flow cytometry staining was performed on single-cell suspensions from each organ. Briefly, cells were first incubated with efluor780-coupled Fixable Viability Dye (Thermo Scientific) for 20 minutes at 4°C to label dead cells. Surface staining was then performed for 45 minutes at 4°C in PBS (TFS) supplemented with 1\% FBS (BioWest) and 0.09\% NaN3 (Sigma-Aldrich). Cells were then fixed and permeabilized in 96 wells plates using 200 uL of BrdU staining solution from the BrdU Staining Kit for Flow Cytometry APC (ThermoScientific) according to manufacturer instructions. The following mAbs(clones) were used: CD8(53.6.7), CD45.1 (A20) from BD Biosciences, CD44(IM7.8.1), Bcl2 (BCL/10C4) and CCL5 (2E9) from Biolegend and Ki67 (SolA15) and CD49d (R1-2) from Thermofischer Scientific. Samples were acquired on a FACS LSR Fortessa (BD biosciences) and analyzed with FlowJo software (TreeStar).

### Estimation of proliferation and differentiation rates of early effector CD8+ T cells

Neglecting CD8+ CD44+ effector cells death over the 24 hours period between BrdU injection and sample collection, we consider that early effector (CD44+ Bcl2-Ki67+, Crauste et al., 2017) CD8 T cells can either proliferate or differentiate. Upon BrdU injection, proliferating cells incorporate BrdU, therefore early effector cell proliferation rate can be approximated by the ratio: #CD44+ BrdU+ cells/ #CD44+ Bcl2-Ki67+ cells (which is equivalent to assuming a linear proliferation rate and that all proliferating cells are BrdU+).

The number of BrdU+ late effector (CD44+ Bcl2-Ki67-, Crauste et al., 2017) cells one day after BrdU injection corresponds to the fraction of BrdU+ early effector cells that have differentiated following BrdU injection. Hence, the differentiation rate of early effector cells into late effector cells is approximated by the ratio: #CD44+ Bcl2-Ki67-BrdU+ cells/ #CD44+ Bcl2-BrdU+ cells.

### In vivo memory precursor cell number calculation

The number of MP present in the spleen of animals at each time point was estimated for each cluster by multiplying the number of cells recovered at this time point (given by the number of cells collected in Yao et al. (2019) by the percentage of cells in the given cluster (given by the TinGa analysis) and the percentage of MP cells among these (given by the GSEA analysis). Then the number of MP was summed for all 8 clusters to yield the number of MP present in the spleen at a given time point.

### Data preprocessing

#### Single-cell RNAseq data preprocessing

Existing single cell data from Yao et al. (2019) were used (GEO, accession no. GSE119943). A feature-barcode matrix by replicate was generated using the Cell Ranger v.3.1 software (10X genomics) and only effector CD8 T cells in acute infection sampled at day 4.5 and day 7 post infection were kept for the analysis. The two replicates were pooled since no batch effect was observed. The cell filtering was made with the scater package (McCarthy et al., 2016). Briefly, cells with a log-library size and a log-transformed number of expressed genes that were more than 3 median absolute deviations below the median value were excluded. The cells with less than 5 % of mitochondrial counts were kept. These criteria were applied separately on the cells from day 4.5 and day 7 leading to 20,295 cells that were kept in total. The data was then normalized using the sctransform function in Seurat (Hafemeister et al., 2019) and variable genes were selected based on variance modelling statistics from the modelGeneVar function in Scran (Lun et al., 2016). The log-normalized expression values of the 2,000 highly variable genes were used for downstream analysis.

To validate our results, a second dataset from Kurd, et al. were used (GEO, accession no. GSE131847). Pre-processed count matrices of cells sampled at day 4, 5, 6, 7 and 10 (replicate 1 only) were pooled (9,614 total cells) and genes detected in less than 1% of the total cells were removed. The data was then normalized using sctransform function and 1,573 highly variable genes were selected by setting the *variable.features.rv.th* parameter to 1.3 (default value).

#### Cell type classification

The cells were automatically annotated and the cell type to which they best corresponded was defined using the SingleR R package (Aran et al., 2019). The labelled normalized expression values of 830 microarray samples of pure mouse immune cells, generated by the Immunologic Genome Project (ImmGen), were used as reference. Cells that were clearly identified as non-T cells (7 B cells, 2 dendritic cells, 3 fibroblasts, 25 macrophages and 62 monocytes) were removed before further analyses were applied.

### Advanced analyses

#### Cell cycle assignement

The Seurat R package was used to classify cells into G1, S or G2/M phases. The classifier relies on a list of genes from Tirosh et al. (2016), that contains markers of the G2/M and S phase. It attributes a class to each cell with a certain probability, with the possibility to attribute the G1 class to cells for which the G2/M or S scores were low.

#### Trajectory inference

Two recently published trajectory inference tools, Slingshot and TinGa, were used to identify a trajectory in the data. The normalized data was first wrapped into a dataset object with the dynwrap R package. The slingshot implementation in dynwrap, as found on the github/dynverse/dynwrap github page, was applied to the data using the default parameters. The TinGa implementation as found on the github/Helena-todd/TInGa repository was applied to the data using the default parameters. The dynplot R package was then used for an easy visualization of the resulting trajectories.

#### Generating heatmaps of gene expression along trajectories

We used the plot\_heatmap() function from the dynplot package to visualize the expression of specific genes along the Slingshot and TinGa trajectories. We either used the function as a discovery tool to identify the top n genes that varied the most along the trajectories, or we provided lists of genes associated with a certain signature to see in which parts of the trajectories these genes were the most expressed.

#### Differential expression analysis

The transitional populations that were identified along the TinGa trajectory were used as clusters defining similar cells. Differential expression analysis was performed between these clusters using the Seurat R package. Wilcoxon rank sum tests were applied and genes were selected as differentially expressed if the difference in the fraction of detection of the gene between the two compared groups of cells was higher than 0.25, and if the log fold-change difference between the two groups was higher than 0.3. The differentially expressed genes were then visualized using the triwise R package (VanLaar et al., 2016) and in a volcano plot that was generated manually in R with the ggplot2 R package.

#### Gene Set Enrichment Analysis

Gene rankings were computed in cells using the AUCell R package. This allowed to identify cells that showed specific gene signatures. Of the 122 genes described as associated with a memory-precursor signature by (Yao et al., 2019), only 42 genes were present in the 2,000 HVGs that we selected. We thus decided to use all genes available instead of restricting ourselves to the 2,000 HVGs for this analysis. 833 cells out of the 20,196 studied acute responding CD8 T cells were assigned to a memory precursor signature.

#### Inferring the number of memory precursors in the spleen

The number of memory precursors in the spleen was calculated based on the percentage of memory precursors identified by gene set enrichment among total day 4.5 or day 7 cells and the average number of CD8 T cells found in the spleen of mice on those same days (Number of MP on day x = \% of MP among single cell from day X * average total number of CD8 T cells in spleen on day X).

#### RNA velocity

Counts of spliced and unspliced abundances were obtained using the Kallisto and Bustools workflow (Melsted et al., 2021). Raw fastq files were pseudo-aligned on Ensembl’s *Mus musculus* reference transcriptome using release 97. Only cells which passed previously described preprocessing steps were kept. To infer RNA velocities and predict cell-specific trajectories, scVelo version 0.2.3 (Bergen et al., 2020) was used. As described in Bergen et al. (2020), velocities were estimated using the dynamical model and the neighborhood graph was computed on the PCA representation using 50 components. The velocity graph was computed with parameter *n_neighbors* set to 20. Other parameters were set to default values. Per-cell MDS coordinates obtained in TinGa were imported into scVelo to project RNA velocities in the same reduced embedding. The 50 genes best fitting scVelo’s model were selected and divided into *Cell-Cycle, Migration* and *Immune Functions* categories according to their function. Finally, figures were obtained by applying the *velocity_embedding_stream* function.

#### Gene regulatory network inference

The BRED R package was used to identify regulatory interactions between a list of transcription factors (that was identified among the 2,000 HVGs using the database in the org.Mm.eg.db R package, and manually curated), and the 2000 target genes. The scaled importances corresponding to these interactions were filtered, and the top 100 interactions corresponding to the 8 populations identified in the TinGa trajectory were selected, resulting in a gene regulatory network containing 800 interactions. A layout of these interactions was then generated using Cytoscape. In the resulting gene regulatory network, we define modules as groups of target genes linked to one central transcription factor.

## Acknowledgment

We thank C. Yao (NIAMS), J. O’shea (NIAMS), P. Schwartzberg (NIAID) and T. Wu (department of Immunology and Microbiology, University of Colorado School of Medicine) for sharing data and their advice. We acknowledge the contribution of SFR BioSciences (UAR3444/CNRS, US8/Inserm, École Normale Supérieure de Lyon, Université de Lyon) and of the CELPHEDIA infrastructure (http://www.celphedia.eu/), especially the center AniRA in Lyon (AniRA-Cytométrie and AniRA-PBES facilities). This work was supported by INSERM, CNRS, Université de Lyon, ENS Lyon, région Auvergne-Rhône-Alpes (Ingerence pack ambition) and ANR (MEMOIRE ANR-18-CE45-0001). Margaux Prieux has a région Auvergne-Rhône-Alpes PhD fellowship.

## Competing interests

The authors declare no competing financial interests.

## Notes

### Competing Interest Statement

The authors have declared no competing interest.

