## Supplementary_materials for "CD8 memory precursor cells generation is a continuous process"

### Supplementary figures

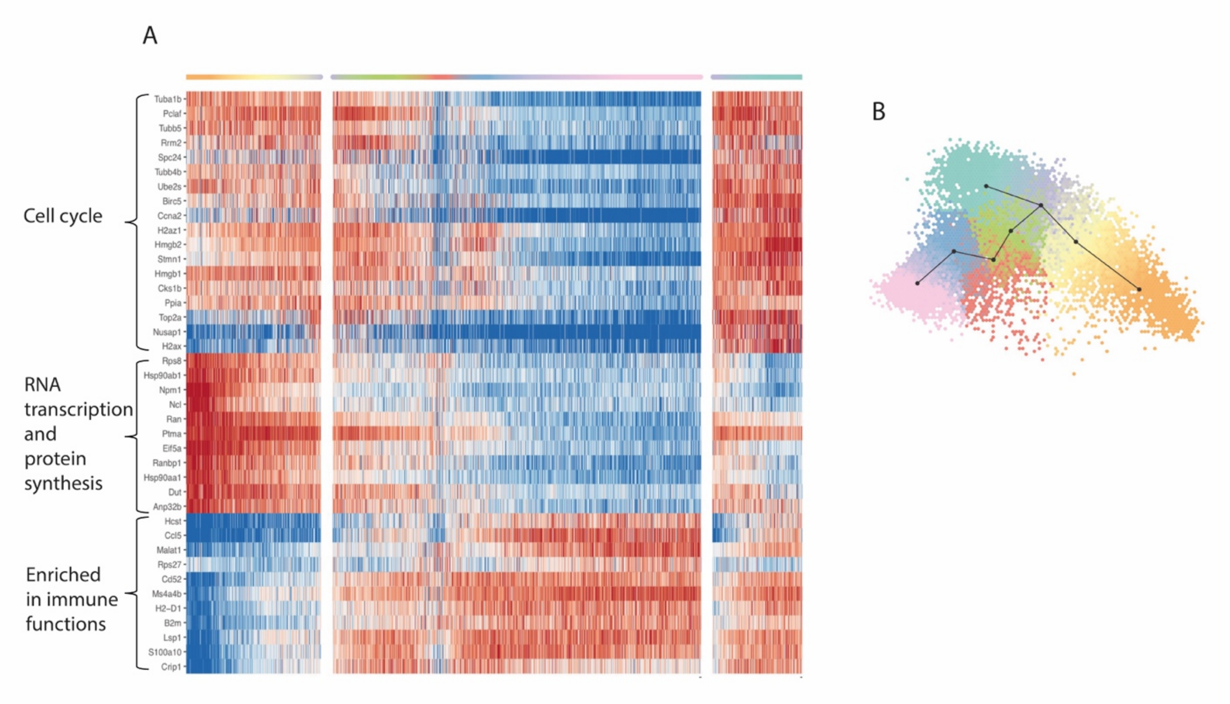


**Supplementary figure 1: (A)** Heatmap of the 40 main genes that were associated with the TinGa trajectory. The genes are represented in rows and the cells are represented in columns and are ordered along the TinGa trajectory. The three branches of the trajectory can be identified with milestone colors on top of the heatmap and numbers at the bottom of the heatmap. Gene expression is depicted from low (blue) to high (red) values in the heatmap. **(B)** Trajectory identified by TinGa when the top 10 percent (= 1300) of the most highly variable genes instead of the 2000 HVGs used in **(A)** and other figures.

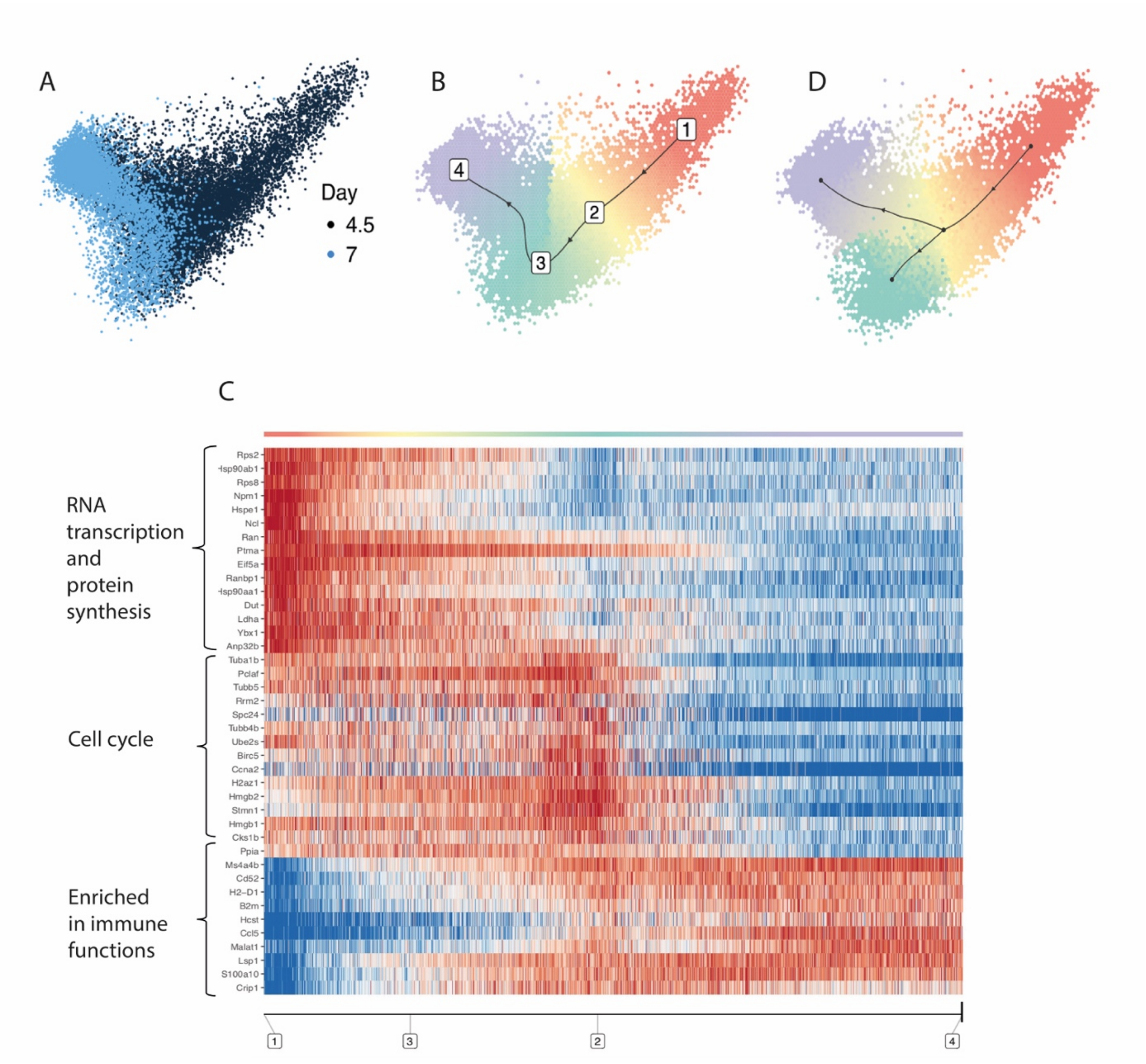


**Supplementary figure 2:** Slingshot trajectory. **(A, B)** Visualisation of the cells in a 2D space computed by principal component analysis. **(A)** The cells were colored according to the two experimental time points 4.5 and 7 days post LCMV-Armstrong infection. **(B)** Slingshot recovered a linear trajectory that is represented by a black directed line. Numbers along the trajectory are used as milestones. **(C)** Heatmap of the 40 main genes that were associated with the Slingshot trajectory. The genes are represented in rows and the cells are represented in columns, re-ordered along the trajectory. The trajectory milestone structure is reported at the bottom of the heatmap as a visual support, and the cluster associated colors are represented at the top of the heatmap. Gene expression is depicted from low (blue) to high (red) values in the heatmap. **(D)** Trajectory identified by Slingshot using the top 10 percent (= 1300) of the most highly variable genes instead of the 2000 HVGs described in this paper.

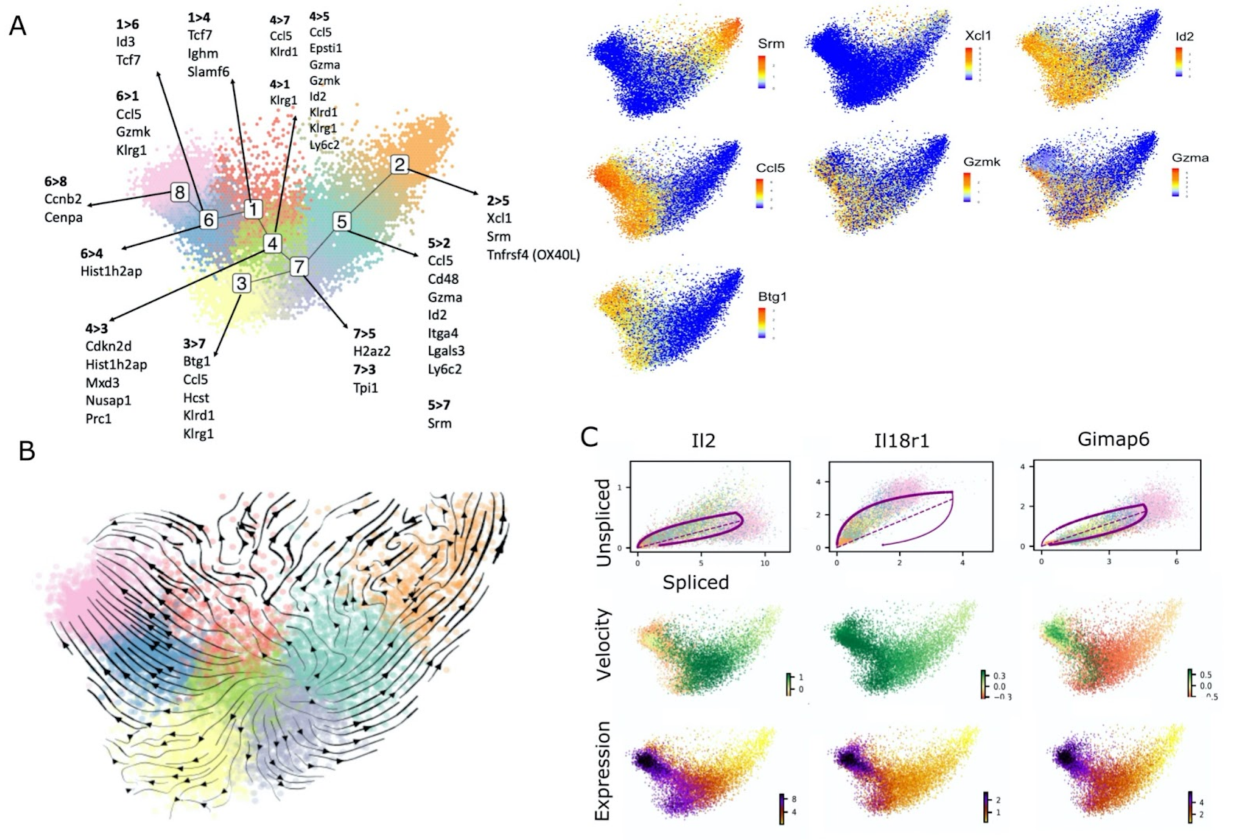


**Supplementary figure 3: (A)** Left panel: Differential expression analysis between two clusters of cells in the same phase of the cell cycle along the TinGa trajectory. Only the genes that are shared between the phases of the cycle were kept. The genes over-represented in paired neighbor-cluster comparative analyses are listed. Right panel: expression of some representative genes was highlighted on the TinGa trajectory. **(B)** Velocities were calculated using the top 50 genes best fitting the dynamical model. **(C)** Illustration of the genetic dynamics of 3 immune function-related genes. Left column: scVelo's dynamical model fitting on the spliced and unspliced counts. Middle column: velocity vectors obtained for each cell on a single gene, positive (green) values indicate cells moving towards regions of higher expression, negative (red) values indicate cells moving towards lower expression. Right column: First moment ('Ms' matrix) of expression levels (spliced counts) for each cell across its nearest neighbors (local mean) for the indicated gene.

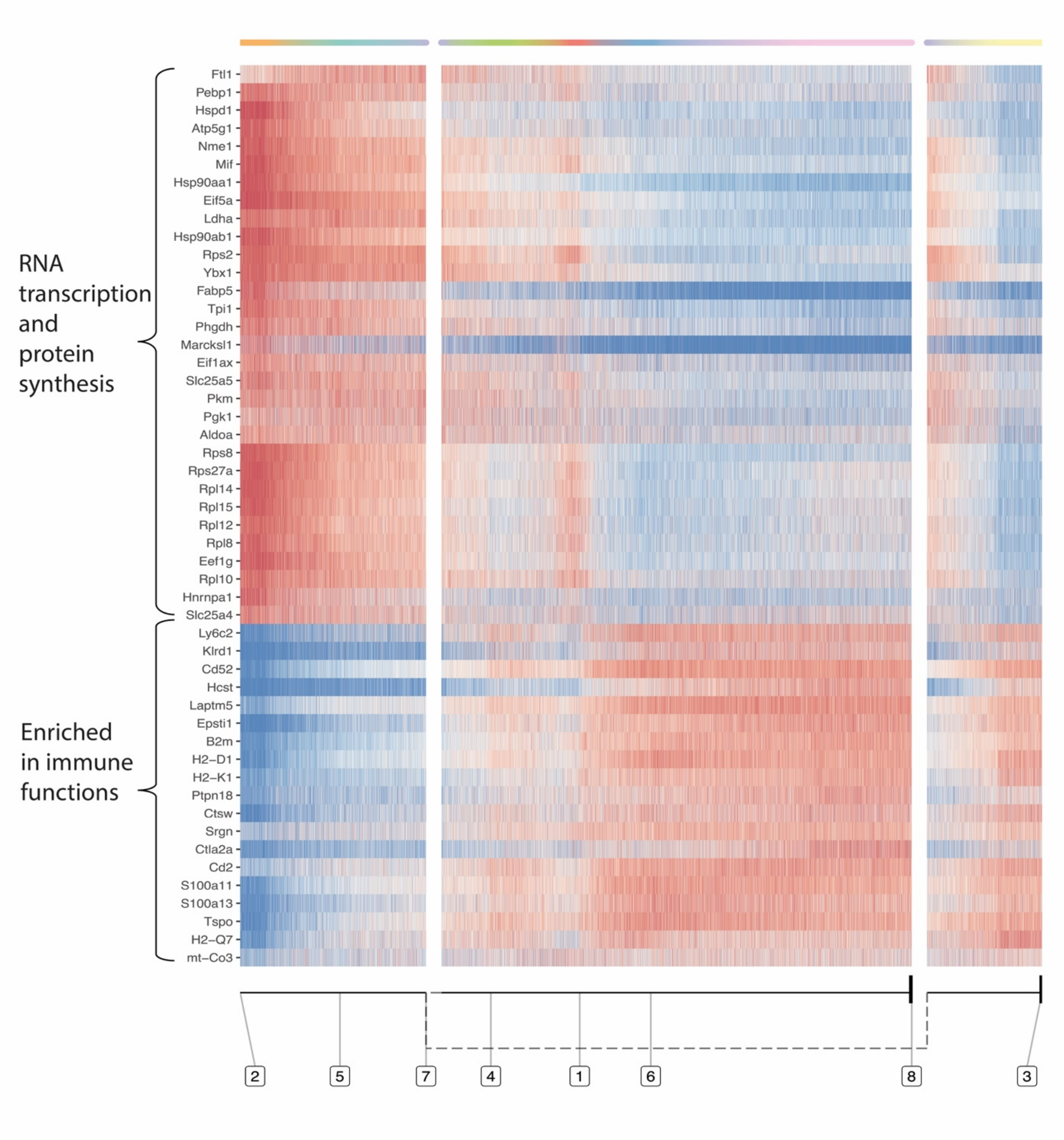


**Supplementary figure 4:** Ybx1 gene module heatmap along TinGa's trajectory represented as explained in the legend to Supplementary figure 1. Two different groups of genes can be identified. Genes at the top of the heatmap are enriched at the beginning of the trajectory and are involved in RNA and protein synthesis. Genes at the bottom of the heatmap are expressed at the end of the trajectory (in clusters 6 and 8) and code for immune receptors.

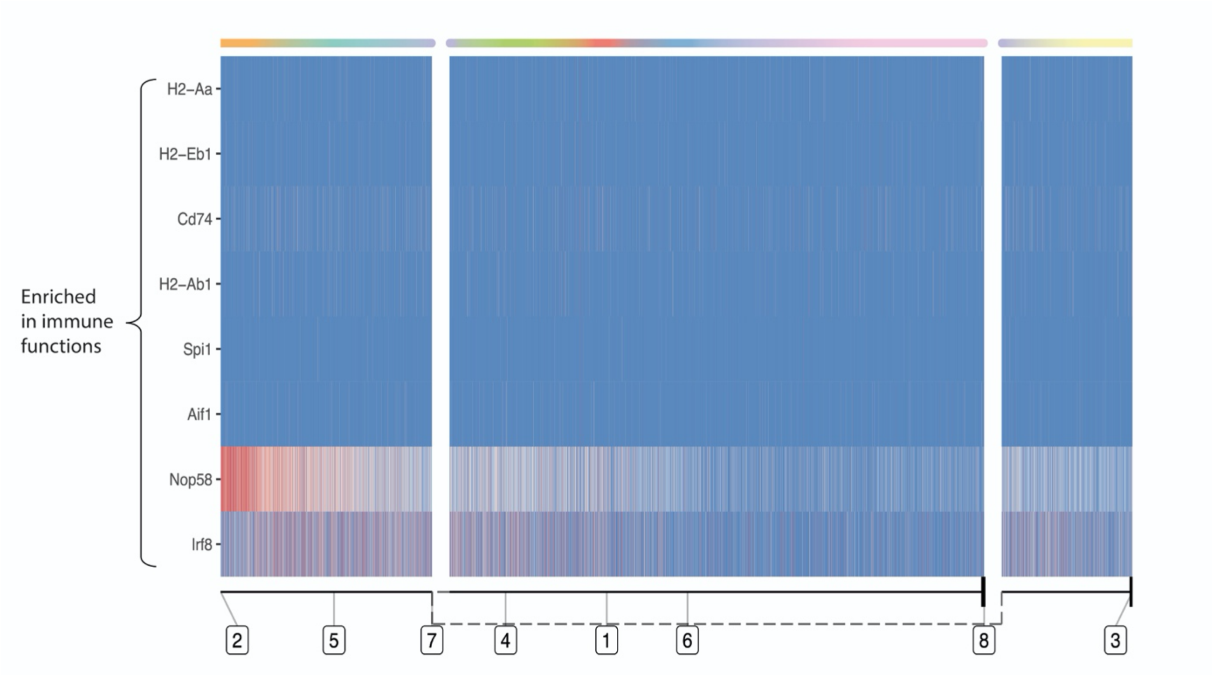


**Supplementary figure 5:** Spi1 gene module heatmap along TinGa's trajectory represented as explained in the legend to Supplementary figure 1.

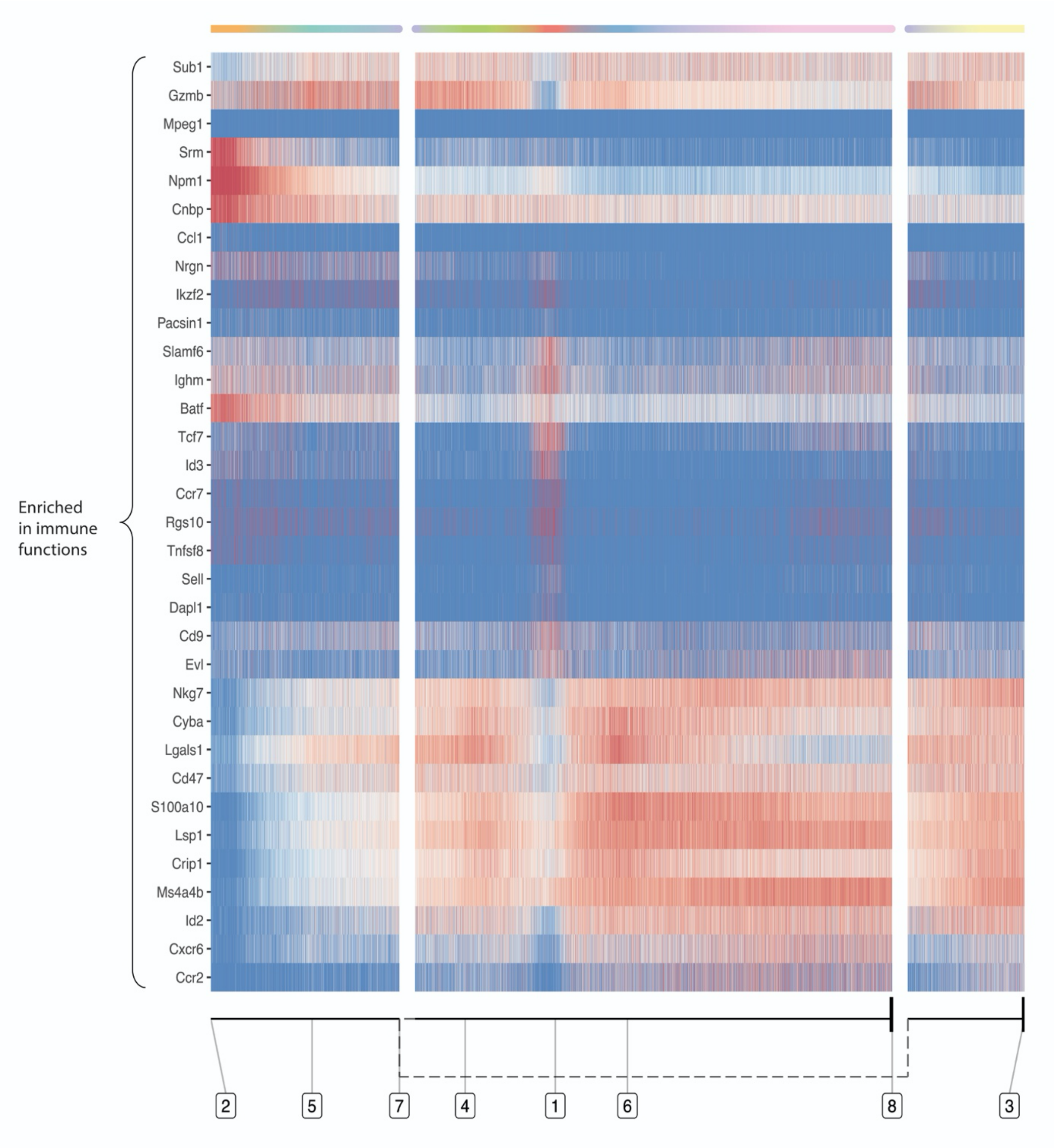


**Supplementary figure 6:** Tcf7/Id2/Phb2 gene module heatmap along TinGa's trajectory represented as explained in the legend to Supplementary figure 1.

A

| Day | G1 | G2/M | S | Total |
|---|---|---|---|---|
| 4.5 | 407 | 296 | 416 | 1119 |
| 7 | 53 | 29 | 60 | 142 |
| Total | 460 | 325 | 476 | 1261 |

B

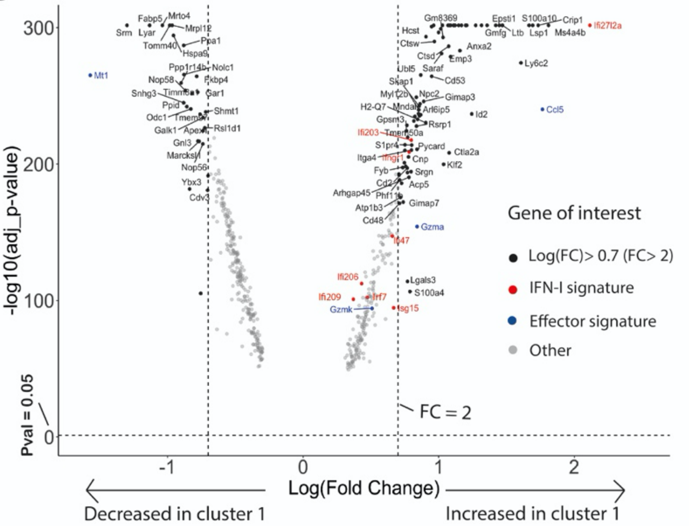


C

| Gene | p_val | avg_logFC | p_val_adj |
|---|---|---|---|
| Tcf7 | 0 | 0.68 | 0 |
| Id3 | 9.62e-220 | 0.54 | 1.92e-216 |
| Ltb | 4.30e-130 | 0.49 | 8.61e-86 |
| Klrg1 | 1.02e-89 | -0.57 | 2.04e-86 |

**Supplementary figure 7:** Analysis of Cluster 1. **(A)** The distribution of the cells from cluster 1 in the cycle phases at each experimental time point. **(B)** Volcano plot of the genes that were differentially expressed between cluster 1 and cluster 2. The genes that change by a fold-change < -2 or > 2 are pointed in black. The genes associated with a type 1 IFN signature are colored in red and the genes that are associated with an effector signature are colored in blue. **(C)** Four genes were differentially expressed in cluster 1 compared to the rest of the cells. The p-values, averaged log fold-changes and the adjusted p-values are reported in this table. Positive values of averaged log fold-changes indicate that the feature is highly expressed in the cluster 1.

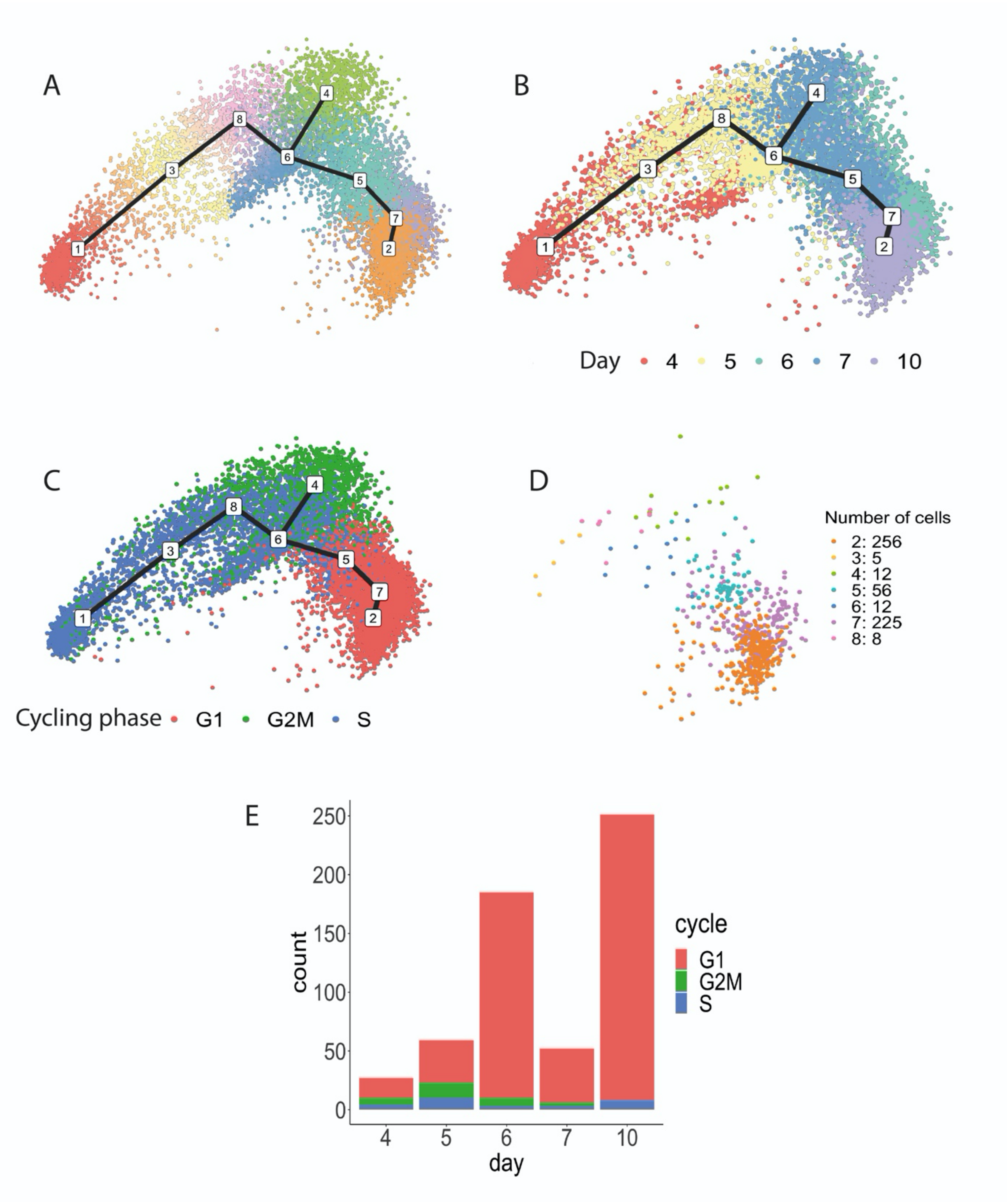


**Supplementary figure 8: (A)** The TinGa algorithm applied on the Kurd et al. dataset identifies a branching trajectory. **(B)** All cells were colored according to their sampling day (4, 5, 6, 7 and 10 post LCMV-Armstrong infection) along the TinGa trajectory. **(C)** All cells were colored according to the phases of the cell cycle along the TinGa trajectory. **(D)** MP cells colored by clusters along the TinGa trajectory. The number of MP cells per cluster is given in the legend. **(E)** MP cell count in cycle phase (G1, S, G2M) across the day post-LCMV infection were represented.

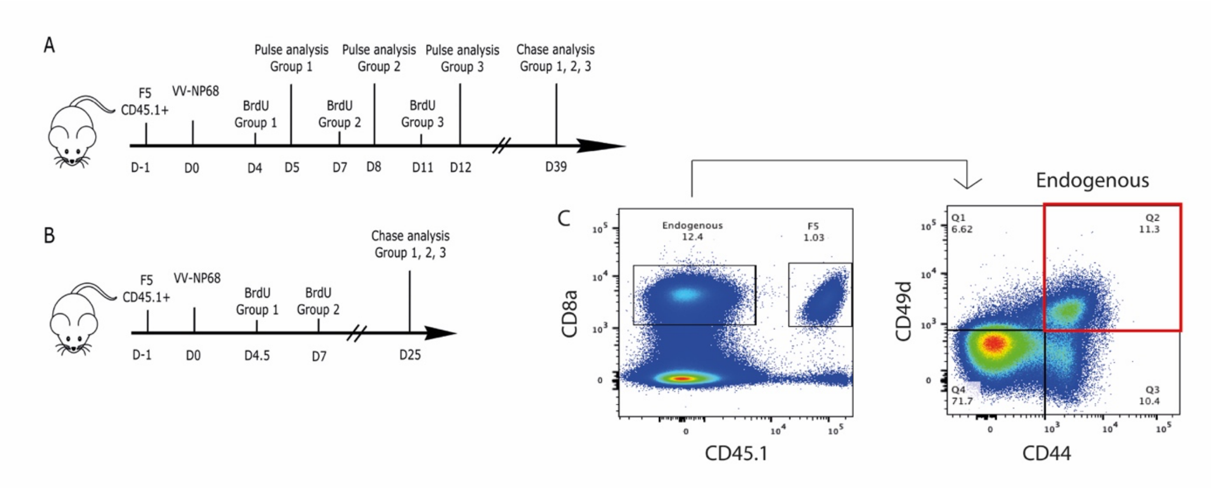


**Supplementary figure 9: (A)** Naive CD45.1 F5 TCR-tg CD8 T cells were transferred to C57BL/6 congenic recipients (n = 5 per group) one day prior i.n. infection with VV-NP68. Mice then received one BrdU injection (2 mg i.p.) on day 4 (Group1), day 7 (Group2) or day 11 (Group3). BrdU labelling was determined by flow cytometry on cells collected 24h after BrdU injection (pulse) or 39 days post infection (chase). **(B)** Naive CD45.1 F5 TCR-tg CD8 T cells were transferred to C57BL/6 congenic recipients (n = 5 per group) one day prior i.n. infection with VV-NP68. Mice then received one BrdU injection (2 mg i.p.) on day 4.5 (Group1) or day 7 (Group2). BrdU labelling was determined by flow cytometry on cells collected 25 days (chase) after infection. **(C)** Gating strategy for the identification of antigen-specific endogenous memory CD8 T cells.

### Supplementary tables

**Supplementary Table** 1: differential expression analysis between two clusters of cells in the same phase of the cell cycle along the TinGa trajectory. Only the genes that are shared between the phases of the cycle were kept.

| gene | p_val | avg_logFC | pct.1 | pct.2 | p_val_adj | comparison |
|---|---|---|---|---|---|---|
| 1500009L16Rik | 1,83E-185 | -0,9245634 | 0,405 | 0,875 | 3,6603E-182 | S_clust_2_vs_5 |
| 1500009L16Rik | 7,3236E-45 | -0,8358373 | 0,426 | 0,832 | 1,46472E-41 | G2M_clust_2_vs_5 |
| AA467197 | 2,3113E-40 | -0,5588995 | 0,187 | 0,564 | 4,62251E-37 | S_clust_1_vs_6 |
| AA467197 | 2,48E-19 | -0,5216 | 0,157 | 0,475 | 4,97E-16 | G2M_clust_1_vs_6 |
| AA467197 | 2,3337E-33 | -0,5589105 | 0,139 | 0,436 | 4,6674E-30 | G1_clust_1_vs_6 |
| Abracl | 1,936E-184 | -0,6519145 | 0,711 | 0,976 | 3,8727E-181 | S_clust_2_vs_5 |
| Abracl | 3,0943E-43 | -0,5346784 | 0,709 | 0,967 | 6,18852E-40 | G2M_clust_2_vs_5 |
| Anxa2 | 0 | -1,1958255 | 0,459 | 0,98 | 0 | S_clust_2_vs_5 |
| Anxa2 | 2,7269E-81 | -1,0162313 | 0,491 | 0,972 | 5,45373E-78 | G2M_clust_2_vs_5 |
| Anxa6 | 1,292E-201 | -0,746947 | 0,411 | 0,898 | 2,5848E-198 | S_clust_2_vs_5 |
| Anxa6 | 1,2107E-45 | -0,5937352 | 0,436 | 0,859 | 2,42136E-42 | G2M_clust_2_vs_5 |
| Arl6ip5 | 6,878E-115 | -0,5175268 | 0,479 | 0,828 | 1,3757E-111 | S_clust_2_vs_5 |
| Arl6ip5 | 1,7935E-31 | -0,5145371 | 0,422 | 0,752 | 3,5871E-28 | G2M_clust_2_vs_5 |
| Btg1 | 3,28E-37 | -0,69769 | 0,201 | 0,7 | 6,55E-34 | S_clust_7_vs_3 |
| Btg1 | 6,425E-238 | -0,8264548 | 0,274 | 0,762 | 1,285E-234 | G2M_clust_7_vs_3 |
| Ccnb2 | 1,019E-295 | 0,78872048 | 0,777 | 0,365 | 2,0375E-292 | G1_clust_6_vs_8 |
| Cenpa | 3,192E-172 | 0,6539074 | 0,796 | 0,534 | 6,3849E-169 | G1_clust_6_vs_8 |
| Ccl5 | 9,3103E-74 | -0,9638835 | 0,055 | 0,354 | 1,86205E-70 | S_clust_2_vs_5 |
| Ccl5 | 1,4015E-18 | -0,8803983 | 0,045 | 0,298 | 2,80295E-15 | G2M_clust_2_vs_5 |
| Ccl5 | 7,05E-75 | -2,85729 | 0,473 | 0,96 | 1,41E-71 | S_clust_7_vs_3 |
| Ccl5 | 1,595E-286 | -2,1074316 | 0,492 | 0,902 | 3,1907E-283 | G2M_clust_7_vs_3 |
| Ccl5 | 5,2372E-65 | -1,6667965 | 0,473 | 0,863 | 1,04744E-61 | S_clust_7_vs_4 |
| Ccl5 | 2,621E-86 | -1,2309854 | 0,492 | 0,934 | 5,24205E-83 | G2M_clust_7_vs_4. |

| | | | | | | |
|---|---|---|---|---|---|---|
| Ccl5 | 1,361E-148 | -2,4206412 | 0,603 | 0,988 | 2,7228E-145 | S_clust_1_vs_6 |
| Ccl5 | 1,48E-90 | -2,41001 | 0,735 | 0,994 | 2,96E-87 | G2M_clust_1_vs_6 |
| Ccl5 | 1,506E-174 | -2,6118703 | 0,73 | 0,994 | 3,0123E-171 | G1_clust_1_vs_6 |
| Ccl5 | 7,994E-121 | 1,93999434 | 0,934 | 0,298 | 1,5989E-117 | G2M_clust_4_vs_5 |
| Ccl5 | 9,845E-304 | 2,00673666 | 0,863 | 0,354 | 1,9691E-300 | S_clust_4_vs_5 |
| Cd48 | 9,492E-226 | -0,8214374 | 0,714 | 0,978 | 1,8984E-222 | S_clust_2_vs_5 |
| Cd48 | 1,8146E-54 | -0,6720766 | 0,713 | 0,968 | 3,62929E-51 | G2M_clust_2_vs_5 |
| Cdkn2d | 1,0934E-81 | -0,6580124 | 0,572 | 0,908 | 2,18688E-78 | G2M_CLUST_3_VS_4 |
| Cdkn2d | 6,8909E-35 | -0,5079278 | 0,495 | 0,787 | 1,37819E-31 | S_CLUST_3_VS_4 |
| Crip1 | 0 | -1,2732093 | 0,683 | 0,999 | 0 | S_clust_2_vs_5 |
| Crip1 | 9,1661E-93 | -1,0942829 | 0,734 | 1 | 1,83323E-89 | G2M_clust_2_vs_5 |
| Ctsd | 1,525E-205 | -0,9144151 | 0,305 | 0,841 | 3,0495E-202 | S_clust_2_vs_5 |
| Ctsd | 1,1768E-44 | -0,7333975 | 0,336 | 0,773 | 2,35361E-41 | G2M_clust_2_vs_5 |
| Ctsw | 4,2573E-50 | 0,5229673 | 0,899 | 0,582 | 8,51468E-47 | G2M_clust_4_vs_5 |
| Ctsw | 3,755E-169 | 0,53874008 | 0,893 | 0,603 | 7,5091E-166 | S_clust_4_vs_5 |
| Emp3 | 4,994E-260 | -0,9913708 | 0,332 | 0,919 | 9,9881E-257 | S_clust_2_vs_5 |
| Emp3 | 2,0129E-64 | -0,8513857 | 0,381 | 0,891 | 4,02574E-61 | G2M_clust_2_vs_5 |
| Epsti1 | 2,25E-147 | -0,650919 | 0,083 | 0,577 | 4,5004E-144 | S_clust_2_vs_5 |
| Epsti1 | 2,0845E-37 | -0,5771634 | 0,069 | 0,507 | 4,16893E-34 | G2M_clust_2_vs_5 |
| Epsti1 | 3,003E-103 | 0,88980637 | 0,976 | 0,507 | 6,0057E-100 | G2M_clust_4_vs_5 |
| Epsti1 | 4,26E-295 | 0,81061416 | 0,945 | 0,577 | 8,5208E-292 | S_clust_4_vs_5 |
| Fabp5 | 4,5695E-79 | -0,7346459 | 0,487 | 0,902 | 9,13897E-76 | G2M_clust_4_vs_5 |
| Fabp5 | 1,311E-213 | -0,6087867 | 0,599 | 0,906 | 2,6226E-210 | S_clust_4_vs_5 |
| Gmfg | 1,128E-217 | -0,826008 | 0,398 | 0,906 | 2,2556E-214 | S_clust_2_vs_5 |
| Gmfg | 1,5299E-52 | -0,7070077 | 0,391 | 0,866 | 3,0597E-49 | G2M_clust_2_vs_5 |
| Gnl3 | 4,689E-153 | 0,58313772 | 0,907 | 0,536 | 9,3775E-150 | S_clust_2_vs_5 |
| Gnl3 | 3,4974E-36 | 0,51692839 | 0,827 | 0,471 | 6,99478E-33 | G2M_clust_2_vs_5 |
| Gzma | 9,591E-115 | -1,6245031 | 0,121 | 0,539 | 1,9182E-111 | S_clust_2_vs_5 |

| | | | | | | |
|---|---|---|---|---|---|---|
| Gzma | 3,442E-32 | -1,5232171 | 0,125 | 0,526 | 6,88392E-29 | G2M_clust_2_vs_5 |
| Gzma | 1,1918E-84 | 1,59653422 | 0,965 | 0,526 | 2,38355E-81 | G2M_clust_4_vs_5 |
| Gzma | 1,002E-184 | 1,41031454 | 0,887 | 0,539 | 2,0041E-181 | S_clust_4_vs_5 |
| Gzmk | 6,6426E-32 | -0,5720413 | 0,447 | 0,714 | 1,32853E-28 | S_clust_1_vs_6 |
| Gzmk | 1,46E-29 | -0,69876 | 0,474 | 0,791 | 2,93E-26 | G2M_clust_1_vs_6 |
| Gzmk | 7,0983E-48 | -0,6531595 | 0,435 | 0,738 | 1,41967E-44 | G1_clust_1_vs_6 |
| Gzmk | 6,8159E-44 | 0,60011157 | 0,729 | 0,36 | 1,36319E-40 | G2M_clust_4_vs_5 |
| Gzmk | 1,5531E-89 | 0,50335221 | 0,712 | 0,438 | 3,10618E-86 | S_clust_4_vs_5 |
| H1f5 | 3,5178E-27 | 0,70369207 | 0,348 | 0,039 | 7,03565E-24 | G2M_CLUST_4_VS_6 |
| H1f5 | 7,786E-98 | 0,75391824 | 0,575 | 0,13 | 1,55721E-94 | S_CLUST_4_VS_6 |
| H2-Q7 | 1,059E-188 | -0,7608977 | 0,596 | 0,953 | 2,1174E-185 | S_clust_2_vs_5 |
| H2-Q7 | 7,6701E-45 | -0,6301942 | 0,63 | 0,94 | 1,53403E-41 | G2M_clust_2_vs_5 |
| H2az2 | 2,2492E-52 | -0,5492847 | 0,608 | 0,906 | 4,49833E-49 | S_clust_5_vs_7 |
| H2az2 | 4,70E-107 | -0,62533 | 0,554 | 0,894 | 9,40E-104 | G2M_clust_5_vs_7 |
| H2az2 | 1,21E-68 | 0,6588974 | 0,927 | 0,554 | 2,42007E-65 | G2M_clust_4_vs_5 |
| H2az2 | 6,511E-209 | 0,5912805 | 0,93 | 0,608 | 1,3022E-205 | S_clust_4_vs_5 |
| Hcst | 7,38E-65 | -1,0769 | 0,423 | 0,941 | 1,48E-61 | S_clust_7_vs_3 |
| Hcst | 3,286E-282 | -0,8936202 | 0,429 | 0,879 | 6,5728E-279 | G2M_clust_7_vs_3 |
| Hcst | 4,3728E-91 | 0,69403379 | 0,755 | 0,181 | 8,74569E-88 | G2M_clust_4_vs_5 |
| Hcst | 2,967E-218 | 0,6561389 | 0,717 | 0,229 | 5,9331E-215 | S_clust_4_vs_5 |
| Hist1h2ap | 5,2973E-26 | 0,74382089 | 0,452 | 0,117 | 1,05947E-22 | G2M_CLUST_4_VS_6 |
| Hist1h2ap | 5,234E-103 | 0,99339733 | 0,613 | 0,166 | 1,0468E-99 | S_CLUST_4_VS_6 |
| Hist1h2ap | 4,2524E-64 | -0,9532653 | 0,452 | 0,781 | 8,5048E-61 | G2M_CLUST_3_VS_4 |
| Hist1h2ap | 1,9486E-70 | -0,8244259 | 0,613 | 0,996 | 3,89718E-67 | S_CLUST_3_VS_4 |
| Id2 | 2,2274E-96 | -0,7869124 | 0,14 | 0,513 | 4,45479E-93 | S_clust_2_vs_5 |
| Id2 | 3,7608E-37 | -0,8960688 | 0,166 | 0,601 | 7,52158E-34 | G2M_clust_2_vs_5 |
| Id2 | 1,5111E-83 | 0,96857233 | 0,969 | 0,601 | 3,02217E-80 | G2M_clust_4_vs_5 |
| Id2 | 9,675E-197 | 0,91671013 | 0,879 | 0,513 | 1,935E-193 | S_clust_4_vs_5 |

| | | | | | | |
|---|---|---|---|---|---|---|
| Id3 | 6,6528E-73 | 0,7720574 | 0,437 | 0,029 | 1,33056E-69 | S_clust_1_vs_6 |
| Id3 | 1,15E-23 | 0,507683 | 0,274 | 0,011 | 2,30E-20 | G2M_clust_1_vs_6 |
| Id3 | 3,215E-128 | 0,77259907 | 0,376 | 0,008 | 6,4296E-125 | G1_clust_1_vs_6 |
| Ifi27l2a | 1,722E-103 | -0,7268999 | 0,242 | 0,666 | 3,4435E-100 | S_clust_2_vs_5 |
| Ifi27l2a | 8,1541E-21 | -0,662013 | 0,298 | 0,615 | 1,63082E-17 | G2M_clust_2_vs_5 |
| Ighm | 6,75E-58 | -0,66228 | 0,411 | 0,725 | 1,35E-54 | S_clust_4_vs_1 |
| Ighm | 2,25E-35 | -0,84282 | 0,336 | 0,708 | 4,51E-32 | G2M_clust_4_vs_1 |
| Itga4 | 1,273E-131 | -0,5953305 | 0,24 | 0,7 | 2,5455E-128 | S_clust_2_vs_5 |
| Itga4 | 1,5437E-41 | -0,6253756 | 0,263 | 0,713 | 3,08749E-38 | G2M_clust_2_vs_5 |
| Itgb7 | 4,035E-224 | -0,8612255 | 0,216 | 0,829 | 8,07E-221 | S_clust_2_vs_5 |
| Itgb7 | 4,01E-56 | -0,7546848 | 0,173 | 0,74 | 8,02001E-53 | G2M_clust_2_vs_5 |
| Itm2b | 1,593E-188 | -0,6823005 | 0,704 | 0,973 | 3,1852E-185 | S_clust_2_vs_5 |
| Itm2b | 6,672E-47 | -0,5740743 | 0,699 | 0,963 | 1,3344E-43 | G2M_clust_2_vs_5 |
| Jak1 | 3,2171E-58 | 0,60116348 | 0,856 | 0,421 | 6,43425E-55 | G2M_clust_4_vs_5 |
| Jak1 | 8,01E-166 | 0,5956979 | 0,826 | 0,479 | 1,602E-162 | S_clust_4_vs_5 |
| Klf2 | 5,042E-143 | -0,7974653 | 0,467 | 0,879 | 1,0084E-139 | S_clust_2_vs_5 |
| Klf2 | 9,8936E-47 | -0,8134536 | 0,471 | 0,864 | 1,97872E-43 | G2M_clust_2_vs_5 |
| Klrd1 | 5,05E-37 | -0,93213 | 0,467 | 0,854 | 1,01E-33 | S_clust_7_vs_3 |
| Klrd1 | 1,167E-178 | -0,7614834 | 0,443 | 0,839 | 2,3338E-175 | G2M_clust_7_vs_3 |
| Klrd1 | 4,9769E-32 | -0,6510008 | 0,467 | 0,747 | 9,95388E-29 | S_clust_7_vs_4 |
| Klrd1 | 5,6012E-45 | -0,5192762 | 0,443 | 0,812 | 1,12024E-41 | G2M_clust_7_vs_4. |
| Klrd1 | 4,9481E-94 | 0,93209529 | 0,812 | 0,213 | 9,89622E-91 | G2M_clust_4_vs_5 |
| Klrd1 | 2,863E-204 | 0,87798679 | 0,747 | 0,288 | 5,7268E-201 | S_clust_4_vs_5 |
| Klrg1 | 1,67E-42 | 0,583865 | 0,436 | 0,099 | 3,34E-39 | S_clust_4_vs_1 |
| Klrg1 | 2,77E-23 | 0,531668 | 0,421 | 0,095 | 5,54E-20 | G2M_clust_4_vs_1 |
| Klrg1 | 2,00E-26 | -0,69734 | 0,157 | 0,549 | 3,99E-23 | S_clust_7_vs_3 |
| Klrg1 | 1,0749E-98 | -0,5549525 | 0,19 | 0,512 | 2,14978E-95 | G2M_clust_7_vs_3 |
| Klrg1 | 1,5089E-62 | -0,8236819 | 0,099 | 0,572 | 3,01777E-59 | S_clust_1_vs_6 |

| | | | | | | |
|---|---|---|---|---|---|---|
| Klrg1 | 3,76E-45 | -0,91912 | 0,095 | 0,612 | 7,52E-42 | G2M_clust_1_vs_6 |
| Klrg1 | 1,5652E-69 | -0,917388 | 0,072 | 0,539 | 3,13041E-66 | G1_clust_1_vs_6 |
| Klrg1 | 8,3743E-39 | 0,55707628 | 0,421 | 0,104 | 1,67486E-35 | G2M_clust_4_vs_5 |
| Klrg1 | 2,8863E-95 | 0,53399858 | 0,436 | 0,141 | 5,77254E-92 | S_clust_4_vs_5 |
| Laptm5 | 7,961E-149 | -0,6065037 | 0,494 | 0,891 | 1,5921E-145 | S_clust_2_vs_5 |
| Laptm5 | 1,3947E-32 | -0,5079673 | 0,505 | 0,832 | 2,78945E-29 | G2M_clust_2_vs_5 |
| Lgals3 | 6,084E-259 | -1,3783836 | 0,385 | 0,928 | 1,2168E-255 | S_clust_2_vs_5 |
| Lgals3 | 2,8358E-63 | -1,1798041 | 0,398 | 0,891 | 5,67154E-60 | G2M_clust_2_vs_5 |
| Lsp1 | 8,843E-300 | -1,1557533 | 0,281 | 0,924 | 1,7687E-296 | S_clust_2_vs_5 |
| Lsp1 | 2,3402E-79 | -1,0174308 | 0,273 | 0,878 | 4,68034E-76 | G2M_clust_2_vs_5 |
| Ly6c2 | 1,7267E-85 | -0,8666487 | 0,191 | 0,563 | 3,45332E-82 | S_clust_2_vs_5 |
| Ly6c2 | 3,5098E-18 | -0,7055798 | 0,215 | 0,507 | 7,01968E-15 | G2M_clust_2_vs_5 |
| Ly6c2 | 2,5392E-81 | 1,08112565 | 0,953 | 0,507 | 5,07849E-78 | G2M_clust_4_vs_5 |
| Ly6c2 | 7,259E-194 | 0,94753361 | 0,907 | 0,563 | 1,4518E-190 | S_clust_4_vs_5 |
| Ms4a4b | 2,117E-289 | -1,1802288 | 0,603 | 0,993 | 4,2331E-286 | S_clust_2_vs_5 |
| Ms4a4b | 2,6039E-81 | -1,0945078 | 0,644 | 0,988 | 5,20788E-78 | G2M_clust_2_vs_5 |
| Ms4a6b | 4,791E-213 | -0,8238616 | 0,618 | 0,974 | 9,5822E-210 | S_clust_2_vs_5 |
| Ms4a6b | 2,9387E-65 | -0,8053935 | 0,616 | 0,957 | 5,87748E-62 | G2M_clust_2_vs_5 |
| Mxd3 | 2,641E-55 | -0,5001029 | 0,369 | 0,764 | 5,28208E-52 | G2M_CLUST_3_VS_4 |
| Mxd3 | 7,1515E-52 | -0,5140897 | 0,354 | 0,791 | 1,4303E-48 | S_CLUST_3_VS_4 |
| Nusap1 | 1,55E-151 | -1,0559958 | 0,384 | 0,943 | 3,0996E-148 | G2M_CLUST_3_VS_4 |
| Nusap1 | 8,166E-123 | -0,9255125 | 0,307 | 0,933 | 1,6332E-119 | S_CLUST_3_VS_4 |
| Pglyrp1 | 1,625E-137 | -0,6186261 | 0,101 | 0,569 | 3,25E-134 | S_clust_2_vs_5 |
| Pglyrp1 | 4,6929E-31 | -0,513608 | 0,093 | 0,485 | 9,3858E-28 | G2M_clust_2_vs_5 |
| Prc1 | 1,096E-101 | -0,7593231 | 0,325 | 0,865 | 2,19192E-98 | G2M_CLUST_3_VS_4 |
| Prc1 | 1,2673E-56 | -0,5685751 | 0,353 | 0,806 | 2,53456E-53 | S_CLUST_3_VS_4 |
| Prr13 | 1,228E-182 | -0,7533671 | 0,347 | 0,836 | 2,4565E-179 | S_clust_2_vs_5 |
| Prr13 | 2,308E-36 | -0,5852993 | 0,37 | 0,755 | 4,61609E-33 | G2M_clust_2_vs_5 |

| | | | | | | |
|---|---|---|---|---|---|---|
| Pycard | 1,31E-242 | -0,9710085 | 0,64 | 0,977 | 2,6202E-239 | S_clust_2_vs_5 |
| Pycard | 5,3463E-58 | -0,8122259 | 0,657 | 0,961 | 1,06927E-54 | G2M_clust_2_vs_5 |
| Rasgrp2 | 6,195E-193 | -0,8072175 | 0,533 | 0,936 | 1,2389E-189 | S_clust_2_vs_5 |
| Rasgrp2 | 1,2827E-43 | -0,6813132 | 0,512 | 0,896 | 2,56535E-40 | G2M_clust_2_vs_5 |
| Reep5 | 1,895E-172 | -0,6061037 | 0,675 | 0,964 | 3,7899E-169 | S_clust_2_vs_5 |
| Reep5 | 1,4153E-42 | -0,5245702 | 0,64 | 0,94 | 2,83057E-39 | G2M_clust_2_vs_5 |
| S100a10 | 0 | -1,2224133 | 0,648 | 0,999 | 0 | S_clust_2_vs_5 |
| S100a10 | 2,9984E-90 | -0,9777056 | 0,72 | 0,997 | 5,99686E-87 | G2M_clust_2_vs_5 |
| S100a11 | 2,601E-200 | -0,8298955 | 0,374 | 0,881 | 5,2025E-197 | S_clust_2_vs_5 |
| S100a11 | 6,5153E-50 | -0,7071857 | 0,408 | 0,861 | 1,30306E-46 | G2M_clust_2_vs_5 |
| S100a13 | 8,2011E-75 | 0,64793529 | 0,953 | 0,645 | 1,64021E-71 | G2M_clust_4_vs_5 |
| S100a13 | 1,356E-203 | 0,56764529 | 0,943 | 0,689 | 2,7114E-200 | S_clust_4_vs_5 |
| S1pr4 | 1,162E-157 | -0,6850742 | 0,213 | 0,717 | 2,3246E-154 | S_clust_2_vs_5 |
| S1pr4 | 8,3464E-42 | -0,6223805 | 0,19 | 0,673 | 1,66928E-38 | G2M_clust_2_vs_5 |
| Selplg | 2,286E-231 | -0,9045231 | 0,42 | 0,903 | 4,5717E-228 | S_clust_2_vs_5 |
| Selplg | 3,7618E-58 | -0,7594169 | 0,401 | 0,86 | 7,52366E-55 | G2M_clust_2_vs_5 |
| Slamf6 | 3,46E-53 | -0,56024 | 0,353 | 0,653 | 6,93E-50 | S_clust_4_vs_1 |
| Slamf6 | 1,58E-31 | -0,6482 | 0,271 | 0,643 | 3,15E-28 | G2M_clust_4_vs_1 |
| Snhg3 | 1,184E-173 | 0,65036631 | 0,923 | 0,63 | 2,3683E-170 | S_clust_2_vs_5 |
| Snhg3 | 4,8113E-44 | 0,54016998 | 0,889 | 0,55 | 9,62259E-41 | G2M_clust_2_vs_5 |
| Sp100 | 1,7462E-74 | 0,66771967 | 0,908 | 0,468 | 3,49244E-71 | G2M_clust_4_vs_5 |
| Sp100 | 1,702E-169 | 0,55067055 | 0,867 | 0,536 | 3,4043E-166 | S_clust_4_vs_5 |
| Srm | 8,802E-259 | 0,99147562 | 0,96 | 0,578 | 1,7605E-255 | S_clust_2_vs_5 |
| Srm | 9,0731E-67 | 0,80595265 | 0,972 | 0,655 | 1,81462E-63 | G2M_clust_2_vs_5 |
| Tcf7 | 1,50E-143 | -0,80465 | 0,059 | 0,506 | 3,00E-140 | S_clust_4_vs_1 |
| Tcf7 | 7,60E-53 | -0,81434 | 0,04 | 0,523 | 1,52E-49 | G2M_clust_4_vs_1 |
| Tcf7 | 3,3458E-77 | 0,81234021 | 0,506 | 0,058 | 6,69162E-74 | S_clust_1_vs_6 |
| Tcf7 | 2,36E-41 | 0,776952 | 0,523 | 0,067 | 4,73E-38 | G2M_clust_1_vs_6 |

| | | | | | | |
|---|---|---|---|---|---|---|
| Tcf7 | 5,516E-179 | 0,99614784 | 0,63 | 0,056 | 1,1032E-175 | G1_clust_1_vs_6 |
| Tnfrsf4 | 9E-158 | 0,74203089 | 0,613 | 0,162 | 1,8001E-154 | S_clust_2_vs_5 |
| Tnfrsf4 | 1,5438E-38 | 0,67534608 | 0,564 | 0,186 | 3,08757E-35 | G2M_clust_2_vs_5 |
| Top2a | 4,463E-28 | 0,79439014 | 0,661 | 0,313 | 8,92599E-25 | G2M_CLUST_4_VS_6 |
| Top2a | 1,8891E-65 | 0,57134516 | 0,795 | 0,544 | 3,77814E-62 | S_CLUST_4_VS_6 |
| Tpi1 | 4,41E-47 | 0,694298 | 0,987 | 0,708 | 8,82E-44 | S_clust_7_vs_3 |
| Tpi1 | 5,863E-231 | 0,6416271 | 0,949 | 0,678 | 1,1727E-227 | G2M_clust_7_vs_3 |
| Tspo | 1,086E-191 | -0,7370557 | 0,24 | 0,811 | 2,1721E-188 | S_clust_2_vs_5 |
| Tspo | 1,809E-49 | -0,6568932 | 0,256 | 0,756 | 3,61791E-46 | G2M_clust_2_vs_5 |
| Tuba1a | 1,796E-190 | -0,7681033 | 0,395 | 0,878 | 3,5913E-187 | S_clust_2_vs_5 |
| Tuba1a | 1,1071E-49 | -0,6546187 | 0,398 | 0,838 | 2,21416E-46 | G2M_clust_2_vs_5 |
| Xcl1 | 8,958E-93 | 0,81585298 | 0,599 | 0,242 | 1,7916E-89 | S_clust_2_vs_5 |
| Xcl1 | 8,8195E-22 | 0,90336876 | 0,595 | 0,294 | 1,76391E-18 | G2M_clust_2_vs_5 |

**Supplementary Table 2:** top 50 genes used for RNA velocity analysis

| | |
|---|---|
| Selenoh | CC |
| Nasp | CC |
| Pola1 | CC |
| Lockd | CC |
| Nap1l1 | CC |
| Cenpp | CC |
| Dut | CC |
| Top2a | CC |
| Rrm2 | CC |
| Gmnn | CC |
| Smc4 | CC |
| Tacc3 | CC |
| Clspn | CC |
| Ckap5 | CC |
| Lmnb1 | CC |
| Pop5 | CC |
| Ddx39 | CC |
| Kif11 | CC |
| Tipin | CC |
| Hsp90ab1 | CC |
| Rnaseh2b | CC |
| Epsti1 | IM |
| Il18r1 | IM |
| Cmtm7 | IM |
| Apbb1ip | IM |
| Skap1 | IM |
| Lck | IM |

| | |
|---|---|
| Park7 | IM |
| Anxa1 | IM |
| Id2 | IM |
| Lgals1 | IM |
| Rnf138 | IM |
| Gimap6 | IM |
| Lsp1 | IM |
| Malat1 | IM |
| Gmfg | MIG |
| Gramd3 | MIG |
| Bin2 | MIG |
| Actb | MIG |
| Cotl1 | MIG |
| Glipr2 | MIG |
| Ppp1r12a | MIG |
| Ezh2 | EPIG |
| Sp100 | EPIG |
| Cbx5 | EPIG |
| 1500009L16Rik | Other |
| Crip1 | other |
| Mrps28 | Other |
| Etfb | Other |
| Gimap4 | apoptosis |

CC: cell cycle; IM: immune; MIG: migration; EPIG: epigenetic

**Supplementary Table 3:** MP gene signature

"Abca3";"Abi2";"Acpp";"Actn1";"Aff3";"Afp";"Akap13";"Als2cl";"Ar";"Arhgap5";"Armcx2";"Axl";"Bach2";"B
cl2";"Btbd11";"Btla";"Ccr6";"Ccr7";"Cd2ap";"Cd55";"Cd7";"Cd86";"Cd9";"Cmpk2";"Cpm";"Ctla4";"Cxcr5"
;"Dapl1";"Ddr1";"Ddx60";"Dhx58";"Dock9";"Elovl6";"Emb";"Evl";"F2rl1";"Faah";"Fam101b";"Fam102a";"
Fam134b";"Fam169b";"Fam26f";"Fam46c";"Fchsd2";"Fcrl1";"Filip1l";"Gpr183";"Hdgfrp3";"Hipk2";"Hvcn
1";"Id3";"Ier3";"Ifitm3";"Ikbke";"Il7r";"Inpp4b";"Ipo4";"Jmjd1c";"Kbtbd11";"Lta";"Ltb";"Ly6e";"Lypd6b";"Ly
smd2";"Mcoln2";"Myc";"Ncoa7";"Nrp1";"Nt5e";"Oas3";"Pacsin1";"Pak6";"Parp12";"Parp8";"Pde4b";"Pdk
1";"Pfn2";"Pgs1";"Pik3Ip1";"Pim2";"Plekha1";"Pou2af1";"Pou6f1";"Ptger2";"Ptpn3";"Qtrtd1";"Rab37";"R
gs10";"Rnf144a";"Rnf213";"Rtp4";"Sell";"Sgms1";"Sh3bp5";"Sirt5";"Slamf6";"Slc11a2";"Smad1";"Socs3
";"Socs5";"Spint2";"Ssbp2";"St6gal1";"St8sia1";"Tacc2";"Taf4b";"Tbc1d4";"Tcf7";"Tlr1";"Tnfrsf25";"Tox";
"Tpd52";"Traf1";"Treml2";"Trib2";"Trmt61a";"Tsc22d1";"Vav3";"Vwa5a";"Wdr12";"Wfikkn2";"Zc3h12d"

**Supplementary Table 4:** list of antibodies tested to characterize BrdU+ memory CD8 T cells generated at day 4 or 7 post-infection.

| Antibody | Source | Identifier | Dilution |
|---|---|---|---|
| PE anti-mouse CCL5 (2E9) | Biolegend | cat# 149103 | 1/400 |
| Biotin anti-mouse Ly6c (AL21) | BD Biosciences | cat# 557359 | 1/800 |
| Biotin anti-mouse Ly108 (13G3-19D) | ebiosciences | cat# 13-1508 | 1/50 |
| BV650 anti-Rat/Mouse CD49a (Ha31/8) | BD Biosciences | cat# 740519 | 1/50 |
| PE-Cy7 anti-mouse CD127 (A7R34) | Ebiosciences | cat# 25-1271 | 1/100 |
| BV510 anti-mouse CD62L (MEL-14) | Biolegend | cat# 104441 | 1/200 |
| FITC anti-mouse KLRG1 (2F1) | Ebiosciences | cat# 11-5893-82 | 1/100 |
| BV421 anti-mouse CXCR3 (CXCR3-173) | Biolegend | cat# 126521 | 1/100 |
